# A DRP-like pseudoenzyme coordinates with MICOS to promote cristae architecture

**DOI:** 10.1101/2023.10.03.560745

**Authors:** Abhishek Kumar, Mehmet Oguz Gok, Kailey N. Nguyen, Michael L. Reese, Jeremy G. Wideman, Sergio A. Muñoz-Gómez, Jonathan R. Friedman

## Abstract

Mitochondrial cristae architecture is crucial for optimal respiratory function of the organelle. Cristae shape is maintained in part by the mitochondrial inner membrane-localized MICOS complex. While MICOS is required for normal cristae morphology, the precise mechanistic role of each of the seven human MICOS subunits, and how the complex coordinates with other cristae shaping factors, has not been fully determined. Here, we examine the MICOS complex in *Schizosaccharomyces pombe*, a minimal model whose genome only encodes for four core subunits. Using an unbiased proteomics approach, we identify a poorly characterized inner mitochondrial membrane protein that interacts with MICOS and is required to maintain cristae morphology, which we name Mmc1. We demonstrate that Mmc1 works in concert with MICOS complexes to promote normal mitochondrial morphology and respiratory function. Bioinformatic analyses reveal that Mmc1 is a distant relative of the Dynamin-Related Protein (DRP) family of GTPases, which are well established to shape and remodel membranes. We find that, like DRPs, Mmc1 self-associates and forms high molecular weight assemblies. Interestingly, however, Mmc1 is a pseudoenzyme that lacks key residues required for GTP binding and hydrolysis, suggesting it does not dynamically remodel membranes. These data are consistent with a model in which Mmc1 stabilizes cristae architecture by acting as a scaffold to support cristae ultrastructure on the matrix side of the inner membrane. Our study reveals a new class of proteins that evolved early in fungal phylogeny and is required for the maintenance of cristae architecture. This highlights the possibility that functionally analogous proteins work with MICOS to establish cristae morphology in metazoans.

## Introduction

Mitochondria are double membrane-bound organelles responsible for myriad cellular functions, including amino acid metabolism, iron-sulfur cluster biogenesis, and energy production via respiration. Efficient respiration relies on the organization of the inner mitochondrial membrane (IMM) into distinct morphological domains, including cristae invaginations and the boundary, an area of close apposition between the IMM and outer mitochondrial membrane (OMM)^1^. The boundary and cristae are separated by cristae junctions, narrow tubular membrane necks that are thought to act as a diffusion barrier between the domains. Numerous factors coordinately give rise to IMM ultrastructure, including the dimerization of ATP synthase, the unique phospholipid composition of the IMM, the IMM fusion machinery, and the Mitochondrial Contact Site and Cristae Organizing System (MICOS) complex^2,3^.

MICOS is a ∼megadalton-sized IMM complex that enriches at cristae junctions and whose loss leads to drastically altered cristae morphology and respiratory dysfunction^4–6^. In accordance with its critical importance for mitochondrial function, the five core MICOS subunits (Mic60, Mic10, Mic19, Mic26, and Mic12) are conserved between the budding yeast *S. cerevisiae* and humans, with additional subunits in each that arose from gene duplications^7,8^. In both organisms, MICOS is organized into two major subcomplexes whose principal components are Mic10 and Mic60, respectively^9^. Mic10 homo-oligomers contribute to membrane tubulation, and accordingly, Mic10 is thought to directly shape the tubular cristae junctions^10,11^. Mic60, along with its partner Mic19, the only non-membrane integral member of the complex, are proposed to stabilize cristae junctions by forming a supportive vault^12^. Additionally, Mic60 and Mic19 interact with the OMM beta-barrel assembly SAM complex^13–16^. This trans-boundary complex between MICOS, SAM, and other OMM proteins, termed the mitochondrial intermembrane space bridging complex (MIB), presents an opportunity for communication between other organelles such as the endoplasmic reticulum and the inside of mitochondria^17^. Other evolutionary newer subunits allow for more complex regulation. These include Mic12 (MIC13 in humans), which is thought to bridge the two subcomplexes, and two apolipoprotein O-like proteins (Mic26 and Mic27) that help coordinate phospholipids, respiratory complexes, and Mic10 function^7,18–22^. However, despite recent advances, the precise role of each subunit and how each protein works together in the context of the holo-MICOS complex has yet to be clearly determined.

Here, we examine the MICOS complex in the fission yeast *Schizosaccharomyces pombe*, which expresses a minimal complement of four MICOS subunits. Mic26 and Mic27, which were independently duplicated in budding yeast and human cells, exist as a single copy in fission yeast. Additionally, Mic12 appears to be absent. Like in budding yeast and human cells, we determine MICOS disruption in *S. pombe* causes altered cristae morphology and respiratory function. Through proteomic analysis, however, we identify a novel MICOS interacting subunit, the poorly characterized mitochondrial protein Mug99. We demonstrate that Mug99 physically and spatially associates with MICOS, working in parallel with the complex to promote cristae morphology. Based on our results, we propose to rename Mug99 as <u>M</u>itochondrial <u>M</u>ICOS interactor required for <u>c</u>ristae morphology 1 (Mmc1). We find that Mmc1 has a predicted structure that is strikingly similar to members of the dynamin related protein (DRP) family of GTPases. Mmc1, however, is a pseudoenzyme that does not contain key residues required for GTP binding or hydrolysis. Like DRPs^23,24^, Mmc1 self-associates in high molecular weight complexes independently of MICOS, suggesting it may work in concert with MICOS to promote cristae stability from the matrix side of the IMM. Phylogenetic analysis reveals that Mmc1 is widespread across fungi and thus represents a new class of proteins regulating cristae morphology with deep roots in the fungal lineage.

## Results

### Proteomic analysis of the S. pombe MICOS complex identifies the poorly characterized protein Mmc1

Four core MICOS subunits are annotated in the fission yeast *S. pombe*: Mic60, Mic19, Mic10, and Mic26. To determine the consequence of the loss of each MICOS subunit, we observed mitochondrial morphology by confocal microscopy of wild type and respective deletion cells expressing the OMM protein Tom20-EGFP^25^. In contrast to wild type cells, which have tubular mitochondrial networks in nearly all cells, the absence of MICOS subunits caused characteristic alterations in mitochondrial morphology (Fig. 1A). As previously observed in budding yeast^26^, mitochondria frequently appeared flattened and lamellar in appearance (Fig. 1A-1B). Additionally, as seen in human cells where MICOS subunits are depleted^27,28^, the mitochondria also exhibit ring-like spherical structures in a significant percentage of cells. Notably, in contrast to both budding yeast and human cells, the loss of each core MICOS subunit affected mitochondrial morphology to a similar extent (Fig. 1B).

**Figure 1.**
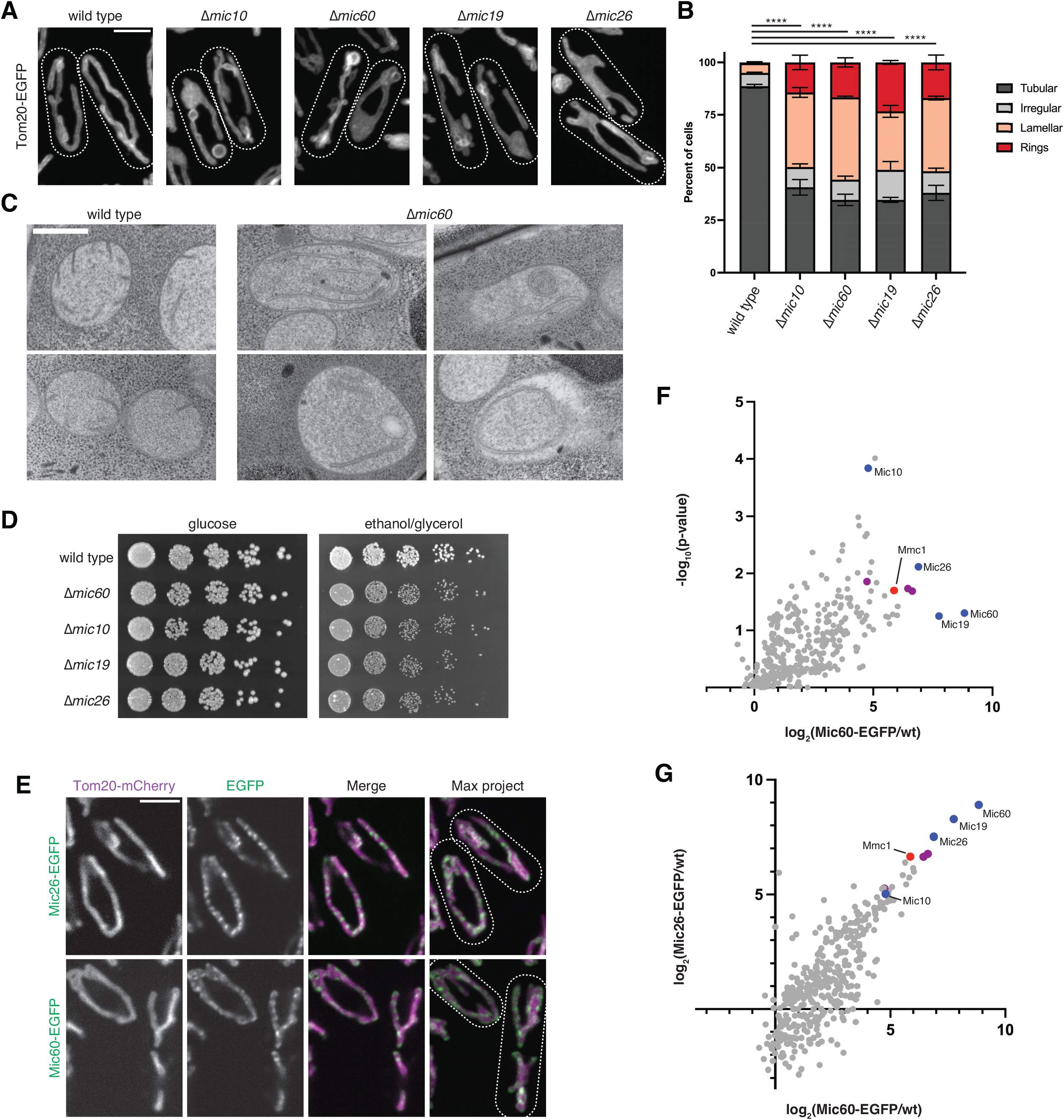
Proteomic analysis of the *S. pombe* MICOS complex identifies the poorly characterized protein Mmc1. **(A)** Maximum intensity projections of confocal microscopy images are shown of the indicated *S. pombe* strains expressing chromosomally tagged Tom20-EGFP and grown to exponential phase in synthetic glucose (EMM) media. **(B)** A graph of the categorization of mitochondrial morphology from cells as in (A). Data shown represent 100 cells per strain in each of three independent experiments, and bars indicate S.E.M. Asterisks (****p<0.0001) represent two-way ANOVA with Tukey’s multiple comparisons test of tubular morphology. **(C)** Representative electron micrographs of the indicated strains grown to exponential phase in EMM media. **(D)** Serial dilutions of the indicated strains grown to exponential phase in YES glucose media and plated on YES media containing glucose or the non-fermentable carbon source ethanol/glycerol and allowed to grow at 30°C for 3 days (glucose) or 10 days (ethanol/glycerol). **(E)** Single plane and, where indicated, maximum intensity projection confocal images of strains co-expressing Mic60 or Mic26 chromosomally tagged with EGFP (green) and Tom20-mCherry (magenta) and grown to exponential phase in YES media. **(F)** A plot of the relative enrichment of total peptide spectral matches of proteins identified by immunoprecipitation and mass spectrometry (IP/MS) analysis of a strain with chromosomally tagged Mic60-EGFP relative to a wild type strain. Data were acquired from three independent experiments. Identified core MICOS subunits are highlighted in blue. MIB proteins (Sam50, Metaxin, DNAJC11) are highlighted in magenta. Mmc1 is highlighted in red. See also Table S1. **(G)** A plot of the relative enrichment of total spectral matches of proteins identified from IP/MS of a Mic60-EGFP or a Mic26-EGFP strain relative to a wild type strain (x-axis and y-axis, respectively). Proteins are highlighted as in (F). Data represent values from three (Mic60-EGFP) or two (Mic26-EGFP) independent experiments. See also Table S2. Scale bars = 4 µm (A, E), 400 nm (C). Cell boundaries are indicated with dotted lines. See also Figures S1 and S2.

To examine mitochondrial ultrastructure in the absence of MICOS, we performed electron microscopy of high-pressure frozen *S. pombe* cells. Wild type cells often had mitochondria with a few cristae present (Fig. 1C). In contrast, Δ*mic60* cells regularly had elongated and stacked cristae membranes, similar to observations in budding yeast^26,29–31^ (Fig. 1C). Consistent with the presence of ring-like structures in fluorescence microscopy images, we also observed membrane rings like those found in MICOS-depleted human cells^18,27,28,32^. Finally, we also observed examples of small, internal membrane-bound and less electron-dense structures within mitochondria as has been seen in MICOS deletion cells in budding yeast^33,34^ (Fig. 1C; see bottom-left Δ*mic60* example).

We next analyzed whether the loss of MICOS subunits impacted the respiratory growth of *S. pombe* cells. In line with previous findings in yeast^26,29,30^ and human cells^35^, disruption of cristae architecture through loss of the core MICOS subunit Mic60 caused a growth defect specifically on media that requires respiration (Fig. 1D). However, unlike in *S. cerevisiae*, where only the absence of Mic10 or Mic60 grossly impacts respiratory growth^26,29,30^, the loss of each core MICOS subunit uniformly affected respiratory growth in *S. pombe* (Fig. 1D).

A common feature of MICOS complex subunits in both budding yeast and human cells is that they exhibit protein stability interdependence. In budding yeast, loss of core MICOS subunits Mic60 and Mic10 leads to the destabilization of their subcomplex partners, Mic19 and Mic27 (a paralog of *Sp*Mic26), respectively^26,29,30^. Variable effects have been observed depending on the human cell line examined, but generally, depletion of human MIC60 leads to severe destabilization of members of both MICOS subcomplexes and disruption of the holo-MICOS complex^28,36,37^. We examined MICOS stability interdependence in *S. pombe* by examining the protein levels of chromosomally HA-tagged proteins in the absence of other MICOS proteins. Each HA-tagged MICOS complex member was functional as assayed by the ability to maintain cell growth on ethanol/glycerol media as compared to the corresponding MICOS deletion (Fig. S1A). In fission yeast, the loss of Mic60 and Mic10 led to destabilization of their partner proteins, Mic19 and Mic26, respectively (Fig. S2). However, Mic19 and Mic26 were more generally destabilized in the absence of all other MICOS subunits (Fig. S2).

Next, we ascertained MICOS complex localization by fluorescence microscopy. We labeled MICOS subunits Mic60 and Mic26 with functional chromosomal EGFP tags (Fig. S1B) and visualized each relative to Tom20-mCherry by confocal microscopy. Both Mic60 and Mic26 appeared in a semi-punctate pattern distributed throughout the mitochondrial network (Fig. 1E). These localization data are similar to observations in both budding yeast and human cells^26,38^, suggesting that MICOS complexes also concentrate at cristae junctions in fission yeast.

While four MICOS complex subunits have been identified in *S. pombe*, no annotated homolog exists for Mic12. Additionally, *S. pombe* only contains a single Mic26 paralog. We considered that additional MICOS subunits may exist, or other proteins may work directly with MICOS in *S. pombe*. We therefore performed immunoprecipitation and mass-spectrometry (IP/MS)-based proteomic analysis of cross-linked lysates from Mic60-EGFP and Mic26-EGFP expressing cells, or a control wild type strain expressing no EGFP tag (Fig. 1F-1G; Supplemental Tables 1 and 2). As expected, all core MICOS subunits were identified among the top hits by both Mic60 and Mic26 (Fig. 1F-1G, blue). The MICOS complex is established to form the MIB complex along with members of the SAM complex^7,13^, and correspondingly, both Sam50 and the SAM subunit Metaxin 1 (Mtx1) were also identified as top interactors (Fig. 1F-1G, magenta). Additionally, the MIB component DNAJC11^7^, while not expressed in budding yeast, has a paralog in *S. pombe* and was also identified by both proteins. Previously observed MICOS-interacting proteins were also identified^26,30,39^, including subunits of the OMM and IMM translocons (TOM and TIM complexes), prohibitins, porin (Por1), the i-AAA protease (Yme1), and OXPHOS proteins such as ATP synthase subunits and NADH dehydrogenase (Nde1). Interestingly, we noticed that one of the top interactors identified by proteomic analysis of both Mic60 and Mic26 was the poorly characterized protein Mmc1 (Fig. 1F-1G, red).

### Mmc1 is a MICOS interactor required for normal cristae morphology

Mmc1, originally named <u>M</u>eiotically <u>u</u>pregulated <u>g</u>ene 99 (Mug99) based on transcriptional profiling^40^, has been shown to localize to mitochondria^41,42^ but its function remains unknown. Mmc1 is 526 amino acids in length and contains a putative N-terminal mitochondrial targeting sequence (MTS) and two consecutive C-terminal hydrophobic segments predicted to be transmembrane domains (Fig. 2A). These signatures indicate that the protein may localize to the IMM with the bulk of the protein facing the mitochondrial matrix (Fig. 2A). However, analyses with InterPro/Pfam^43^ failed to identify any domains indicative of the protein’s function.

**Figure 2.**
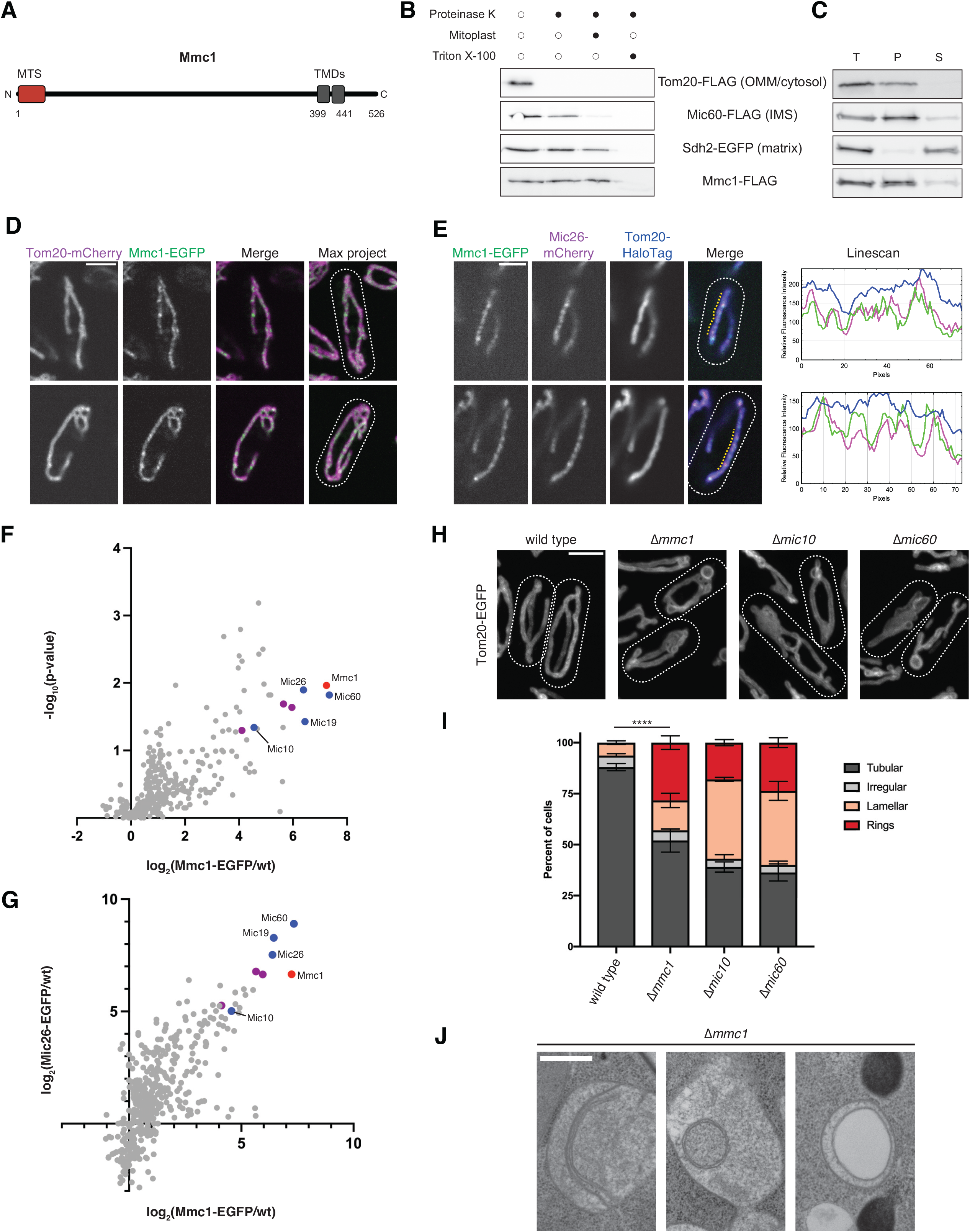
Mmc1 is a MICOS interactor required for normal cristae morphology. **(A)** A schematic of the domain organization of Mmc1. The mitochondrial targeting sequence (MTS) is shown in red and transmembrane domains (TMDs) are shown in gray. **(B)** Western analysis of protease protection assays simultaneously performed on mitochondria isolated from strains expressing the indicated chromosomally tagged proteins. Mitochondria were treated as indicated with proteinase K, mitoplasting (disruption of the outer membrane by a combination of osmotic swelling and mechanical disruption), and/or Triton X-100. **(C)** As in (B) for purified mitochondria that were subjected to alkaline extraction by incubation with 0.1 M Na_2_CO_3_ (pH 11.0). Total (T), pellet (P), and supernatant (S) fractions were collected after ultracentrifugation. **(D)** Single plane (left) and maximum intensity projection (right) confocal images of a strain co-expressing chromosomally tagged Mmc1-EGFP (green) and Tom20-mCherry (magenta) and grown to exponential phase in YES media. **(E)** Single plane confocal microscopy images of a strain co-expressing chromosomally tagged Mmc1-EGFP (green), Mic26-mCherry (magenta), and Tom20-HaloTag labeled with JF646 (blue), and grown to exponential phase in YES media. Fluorescence intensity linescans shown at right correspond to the yellow dotted line shown on the corresponding image. **(F)** A plot of the relative enrichment of total peptide spectral matches of proteins identified by IP/MS analysis of a strain with chromosomally tagged Mmc1-EGFP relative to a wild type strain. Data were acquired from three independent experiments. Identified core MICOS subunits are highlighted in blue, MIB subunits are highlighted in magenta, and Mmc1 is highlighted in red. See also Table S3. **(G)** A plot of the relative enrichment of total spectral matches of proteins identified from IP/MS of strains expressing Mmc1-EGFP or Mic26-EGFP relative to a wild type strain. Proteins are highlighted as in (F). Data represent values from three (Mmc1-EGFP) or two (Mic26-EGFP) independent experiments. See also Table S4. **(H)** Maximum intensity projection confocal images of the indicated strains expressing chromosomally tagged Tom20-EGFP and grown to exponential phase in EMM media. **(I)** A graph of the categorization of mitochondrial morphology from cells as in (H). Data shown represent 100 cells per strain in each of three independent experiments, and bars indicate S.E.M. Asterisks (****p<0.0001) represent two-way ANOVA with Tukey’s multiple comparisons test of tubular morphology. **(J)** Representative electron micrographs of Δ*mmc1* cells grown to exponential phase in EMM media. Scale bars = 4 µm (D, H), 3 µm (E), 400 nm (J). Cell boundaries are indicated with dotted lines. See also Figures S1 and S3.

To determine the localization and topology of Mmc1 we generated a functional chromosomal FLAG-tagged Mmc1 (Fig. S1C) as well as chromosomally tagged versions of other well-characterized mitochondrial proteins. We then purified mitochondria from each respective *S. pombe* strain and assessed protease accessibility. Mmc1-FLAG was detected in the purified mitochondria by Western analysis and was protected from proteinase K digestion, suggesting the protein indeed localizes to mitochondria (Fig. 2B). However, in mitoplasts (mitochondria whose OMM was disrupted by osmotic swelling and mechanical disruption), Mic60-FLAG was selectively degraded while Mmc1-FLAG and the matrix protein Sdh2-EGFP remained intact. Instead, Mmc1-FLAG was only degraded after the combined addition of proteinase K and the detergent Triton X-100, suggesting that the C-terminus of Mmc1 localizes to the matrix. Next, we subjected the purified mitochondria to an alkaline solution of sodium carbonate, which selectively dissociates non-membrane integral proteins from membranes. After ultracentrifugation, Mmc1 was found in the pellet fraction, indicating that it is an integral membrane protein (Fig. 2C). Together, these data are consistent with Mmc1 localizing to the mitochondrial IMM, with both its N- and C-termini exposed to the matrix and likely only a short region (less than 5 amino acids based on bioinformatic predictions) exposed to the intermembrane space (IMS).

We next examined the sub-mitochondrial localization of Mmc1 by confocal microscopy. We generated a functional C-terminal chromosomal EGFP tag of Mmc1 and tested its localization relative to the mitochondrial marker Tom20-mCherry (Fig. 2D and Fig. S1C). As in the case of the MICOS proteins Mic60 and Mic26, Mmc1 was non-uniformly distributed throughout the mitochondrial membrane and had a semi-punctate appearance. To determine whether the distribution of Mmc1 matched that of the MICOS complex, we co-expressed Mmc1-EGFP, Mic26-mCherry, and Tom20-HaloTag in cells. The distribution of Mmc1 appeared similar to Mic26 and the two proteins enriched at the same focal structures (Fig. 2E, see linescan). We next performed IP/MS analysis with anti-GFP conjugated beads from cross-linked lysate from Mmc1-EGFP expressing cells compared to a non-EGFP expressing control. Proteomic analysis of Mmc1-EGFP readily identified all MICOS proteins and several MIB members among the most abundant hits (Fig. 2F, Supplemental Table 3), and the interaction profiles for Mmc1-EGFP and Mic26-EGFP appeared very similar (Fig. 2G, Supplemental Table 4). These data, in combination with previous immuno-EM labeling experiments of MICOS subunits^31,38^, suggest that Mmc1 concentrates at cristae junctions, where it associates in proximity with the MICOS complex.

We then tested whether the loss of Mmc1 affected mitochondrial morphology similarly to the disruption of the MICOS complex. We expressed chromosomally-tagged Tom20-EGFP in Δ*mmc1* cells and visualized mitochondria by confocal microscopy. Approximately half of the Δ*mmc1* cells had abnormal mitochondrial morphology, and many cells contained the lamellar and ring-shaped mitochondria characteristic of cells with MICOS subunit deletions (Fig. 2H-2I). However, the effect of the loss of Mmc1 was slightly less severe, and accordingly, Δ*mmc1* cells did not exhibit an appreciable growth defect on respiratory-requiring media (Fig. S3). Additionally, while lamellar mitochondria are the prominent morphology found in MICOS deletion cells, mitochondrial rings were more prevalent in the absence of Mmc1 (Fig. 2I). We also visualized the mitochondrial ultrastructure of Δ*mmc1* cells by electron microscopy, and consistent with fluorescence microscopy analysis, we could often observe stacked cristae membranes, ringed structures, and less electron-dense internal structures as we observed in Δ*mic60* cells (Fig. 2J). Together, these data indicate that Mmc1 is a MICOS complex-associated protein required for normal cristae morphology.

### Mmc1 associates with MICOS via the Mic10/Mic26 subcomplex

To determine if Mmc1 was affected by disruption of the MICOS complex, we examined its localization in the absence of each core MICOS subunit. We co-expressed Mmc1-EGFP and Tom20-mCherry in either wild type cells or individual MICOS subunit deletions, which were imaged by confocal microscopy. Notably, the semi-punctate Mmc1 distribution we observed in wild type cells was altered in each MICOS subunit deletion. In the absence of Mic60 and Mic19, which together comprise one MICOS subcomplex, Mmc1 concentrated at foci that were more discrete in appearance in the majority of cells (Fig. 3A-B). In contrast, in the absence of Mic10 and Mic26, which are constituents of the second MICOS subcomplex, Mmc1 lost its semi-punctate appearance and was more uniformly localized throughout the mitochondrial network (Fig. 3A-3B).

**Figure 3.**
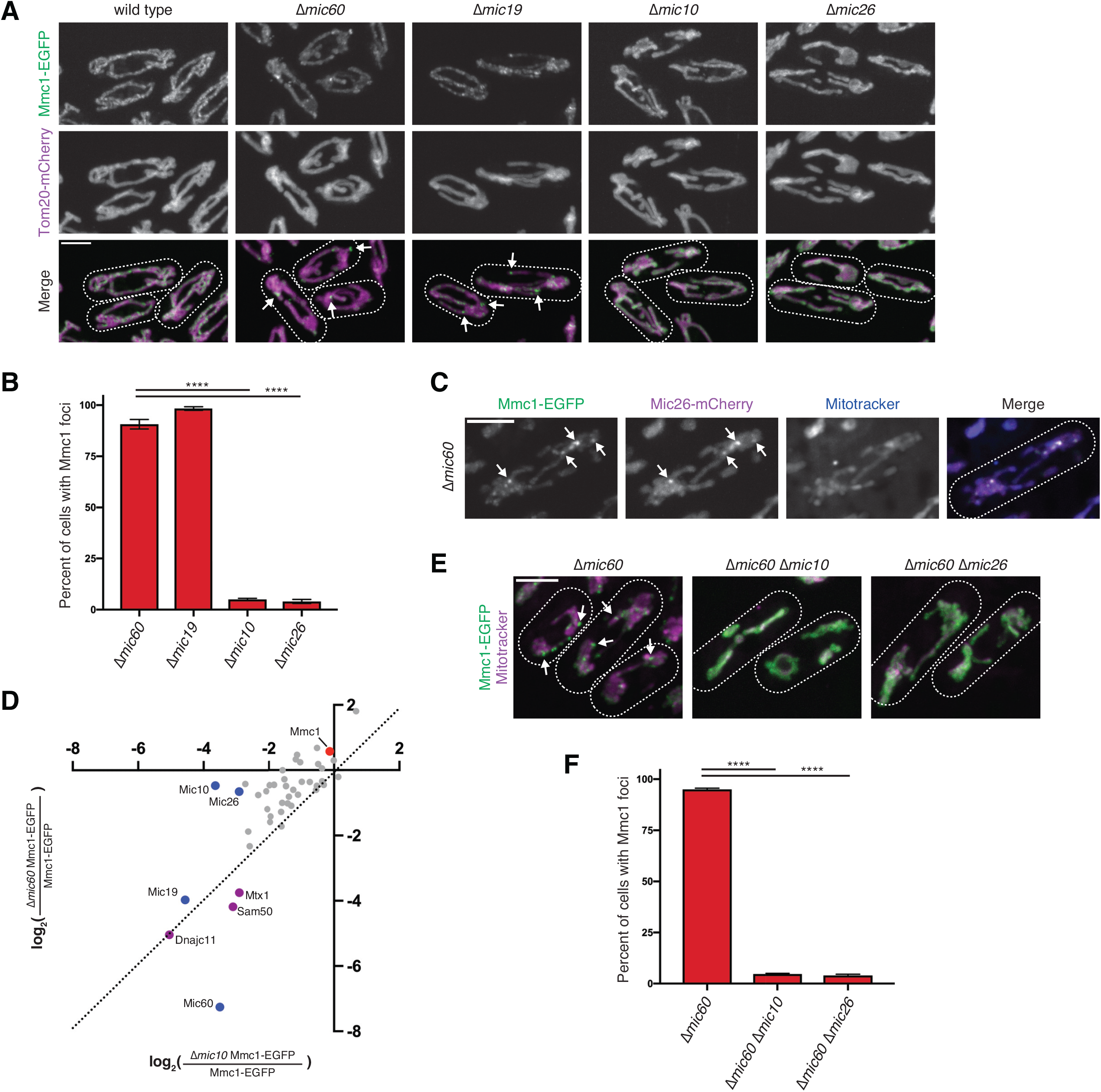
Mmc1 associates with MICOS via the Mic10/Mic26 subcomplex. **(A)** Maximum intensity projection confocal images of the indicated strains co-expressing chromosomally tagged Mmc1-EGFP and Tom20-mCherry and grown to exponential phase in YES media. Arrows mark sites of Mmc1 enrichment in discrete foci. **(B)** A graph depicting the percent of the indicated cells grown as in (A) where Mmc1 localized to discrete foci. Data shown represent 100 cells per strain in each of three independent experiments, and bars indicate S.E.M. Asterisks (****p<0.0001) represent ordinary one-way ANOVA with Dunnett’s multiple comparisons test. **(C)** As in (A) for Δ*mic60* cells co-expressing chromosomally tagged Mmc1-EGFP (green) and Mic26-mCherry (magenta) and stained with Mitotracker Deep Red (blue). Arrows mark sites of Mmc1 and Mic26 co-enrichment. **(D)** A plot of the relative enrichment of total spectral matches of proteins identified from IP/MS of Mmc1-EGFP from Δ*mic10* relative to wild type cells (x-axis) and Δ*mic60* relative to wild type cells (y-axis). All enrichments were computed relative to protein identification in a non-EGFP expressing strain. Identified core MICOS subunits are highlighted in blue, MIB subunits are highlighted in magenta, and Mmc1 is highlighted in red. A dotted diagonal line is shown for reference. Data represent values from two independent experiments. See also Table S5. **(E)** As in (A) for the indicated strains expressing chromosomally tagged Mmc1-EGFP (green) and stained with Mitotracker Far Red (magenta). **(F)** As in (B) for cells grown as in (E). Scale bars = 4 µm (A, C, E). Cell boundaries are indicated with dotted lines.

In budding yeast, Mic27 (a homolog of *Sp*Mic26) localizes to discrete foci in the absence of Mic60 or Mic19^34,44^. We thus asked whether the foci formed by Mmc1-EGFP in Δ*mic60* and Δ*mic19* cells were associated with Mic26. In Δ*mic60* cells co-expressing Mmc1-EGFP and Mic26-mCherry and labeled with Mitotracker, we frequently observed the two proteins concentrating together at the same sites (Fig. 3C). To determine if these focal sites represent an association between Mmc1 and the Mic10/Mic26 subcomplex, we performed IP/MS proteomic analysis of Mmc1-EGFP in wild type cells or in the absence of Mic10 or Mic60. Strikingly, in the absence of Mic10 relative to wild type cells, MICOS and MIB complex members were not readily identified by Mmc1 in IP/MS analysis (Fig. 3D, Supplemental Table 5). However, in Δ*mic60* cells, Mmc1-EGFP retained its ability to associate with both Mic10 and Mic26, but not Mic19 or the MIB complex (Fig. 3D). We then asked whether the Mmc1-EGFP foci detected in Δ*mic60* cells were dependent on the presence of either Mic10 or Mic26. Relative to Mitotracker, Mmc1-EGFP uniformly decorated mitochondria in Δ*mic60 Δmic10 or Δmic60 Δmic26* cells, losing its focal appearance compared to Δ*mic60* cells (Fig. 3E-3F). Together, these data suggest that Mmc1 associates with MICOS via an association with the Mic10/Mic26 subcomplex.

### Exogenous Mmc1 expression in human cells causes the MICOS complex to redistribute

We next sought to ascertain the consequence of Mmc1 expression in a heterologous system to determine its effect on the MICOS complex. Mmc1 is not found in metazoans, and we therefore transiently co-transfected U2OS cells, an immortalized human osteosarcoma cell line, with expression plasmids for Mmc1-GFP and the mitochondrial matrix marker mito-mCherry. Mmc1-GFP localized to mitochondria where it concentrated in stable, discrete puncta distributed throughout the mitochondrial network, particularly in cells with high expression levels (Fig. 4A). To analyze whether these Mmc1 foci were spatially linked to MICOS complexes, we fixed Mmc1-GFP transfected cells and immunolabeled with antibodies against MIC60 and the matrix marker HSP60. In non-transfected cells, MIC60 localized in a semi-punctate pattern relative to HSP60, as expected^27,38^ (Fig. 4B). Remarkably, in cells with high Mmc1-GFP expression, MIC60 often enriched at the sites of Mmc1 foci (Fig. 4B, see arrows and linescan). To ask if this effect was specific to MIC60, we also labeled Mmc1-GFP expressing cells for MIC27 and again could observe that Mmc1 caused redistribution of the MICOS subunit to sites of Mmc1 foci (Fig. 4C). To rule out that exogenous Mmc1-GFP expression caused non-specific protein aggregation, we also examined the relative localization of Mmc1-GFP foci to the IMM protein TIMM23, finding that the protein was not co-enriched at Mmc1-GFP focal sites (Fig. 4D). We also compared Mmc1-GFP foci positioning relative to mtDNA, which resides in nucleoid structures that are localized in the mitochondrial matrix and appear as discrete puncta (Fig. S4). However, Mmc1 and mtDNA localized to distinct punctate sites throughout the mitochondrial network and did not commonly associate in proximity (Fig. S4). Together with our analysis in *S. pombe*, these data additionally indicate that Mmc1 is capable of co-localizing with and influencing the sub-mitochondrial distribution of MICOS subunits in human cells.

**Figure 4.**
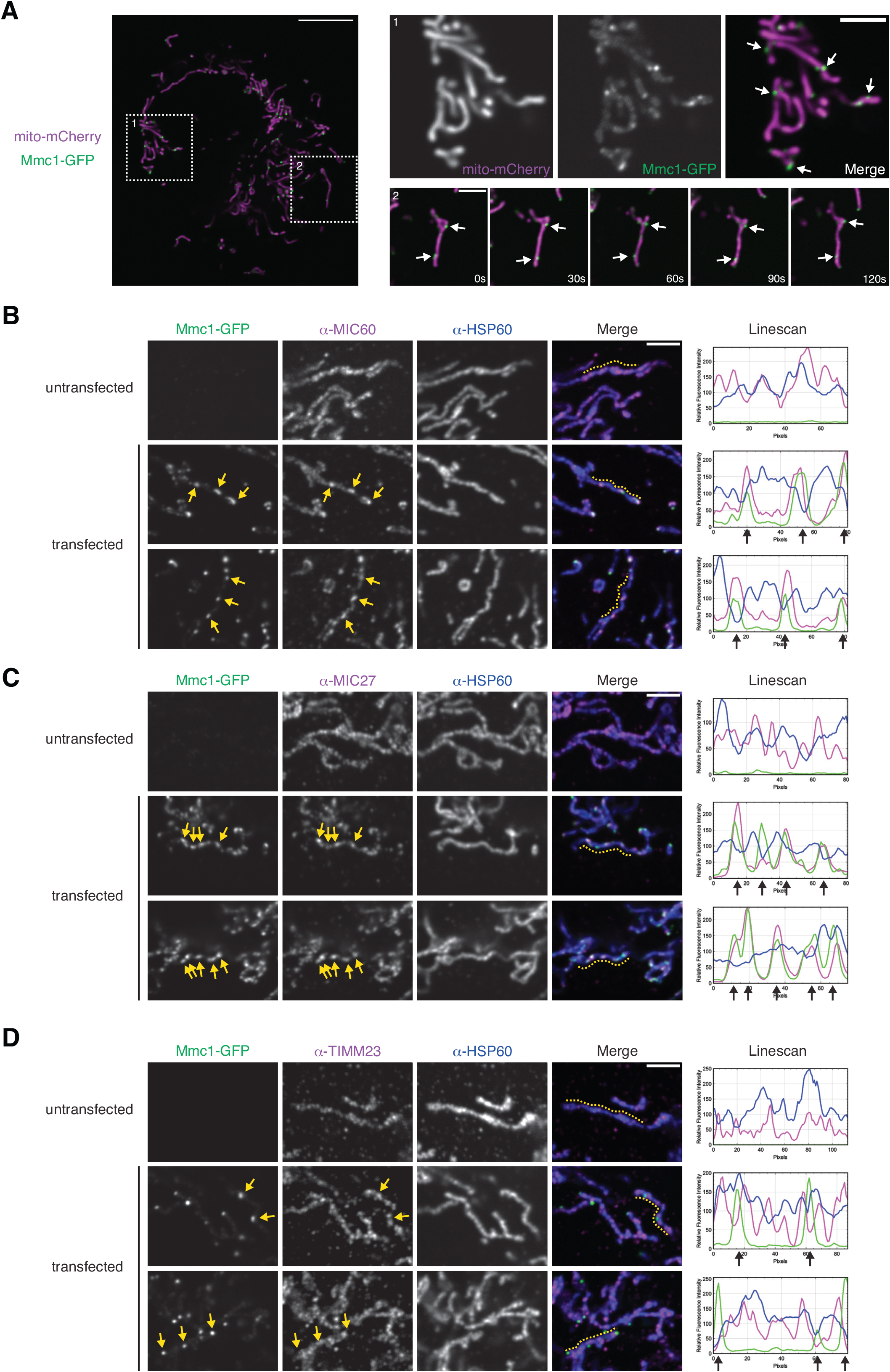
Exogenous Mmc1 expression in human cells causes the MICOS complex to redistribute. **(A)** Left – a single plane confocal microscopy image is shown of a U2OS cell transiently co-transfected with Mmc1-GFP (green) and mito-mCherry (magenta). Dashed box 1 corresponds to the enlargement shown at the top-right. Dashed boxed 2 corresponds to time-lapse microscopy shown at the bottom-right for the indicated intervals in seconds (s). Arrows mark sites of enrichment of Mmc1 in discrete foci. **(B)** Single plane confocal fluorescence microscopy images are shown of a non-transfected U2OS cell (top) or cells transfected with Mmc1-GFP (green). Cells were fixed and immunolabeled with anti-MIC60 (magenta) and anti-HSP60 (blue). Fluorescence intensity linescans shown at right were performed corresponding to the dashed yellow line shown on the merge image. Yellow arrows mark Mmc1 focal enrichments that correspond to the indicated location on the MIC60 image and the associated black arrows marked on linescans. **(C)** As in (B) for cells immunolabeled for MIC27 (magenta). **(D)** As in (B) for cells immunolabeled for TIMM23 (magenta). Note that sites of Mmc1 enrichment do not directly correspond to discrete enrichment of TIMM23 signal. Scale bars = 10 µm (A, left), 3µm (A, right; B-D) See also Figure S4.

### Mmc1 is not required for assembly of the MICOS complex

Given that Mmc1 associates with the MICOS complex, influences MICOS distribution in a heterologous system, and its loss partially phenocopies disruption of MICOS in fission yeast, we hypothesized that Mmc1 may play a role in MICOS assembly. As MICOS proteins exhibit protein stability interdependence (Fig. S2), we asked whether MICOS steady-state levels were affected by the loss of Mmc1 in *S. pombe*. However, the levels of each core HA-tagged MICOS subunit were not substantially altered in the absence of Mmc1 (Fig. 5A). Similarly, Mmc1-FLAG levels were largely unaffected by the loss of individual MICOS subunits, or in Δ*mic10 Δmic60* cells where the MICOS complex is likely destabilized (Fig. S2, Fig. S5).

**Figure 5.**
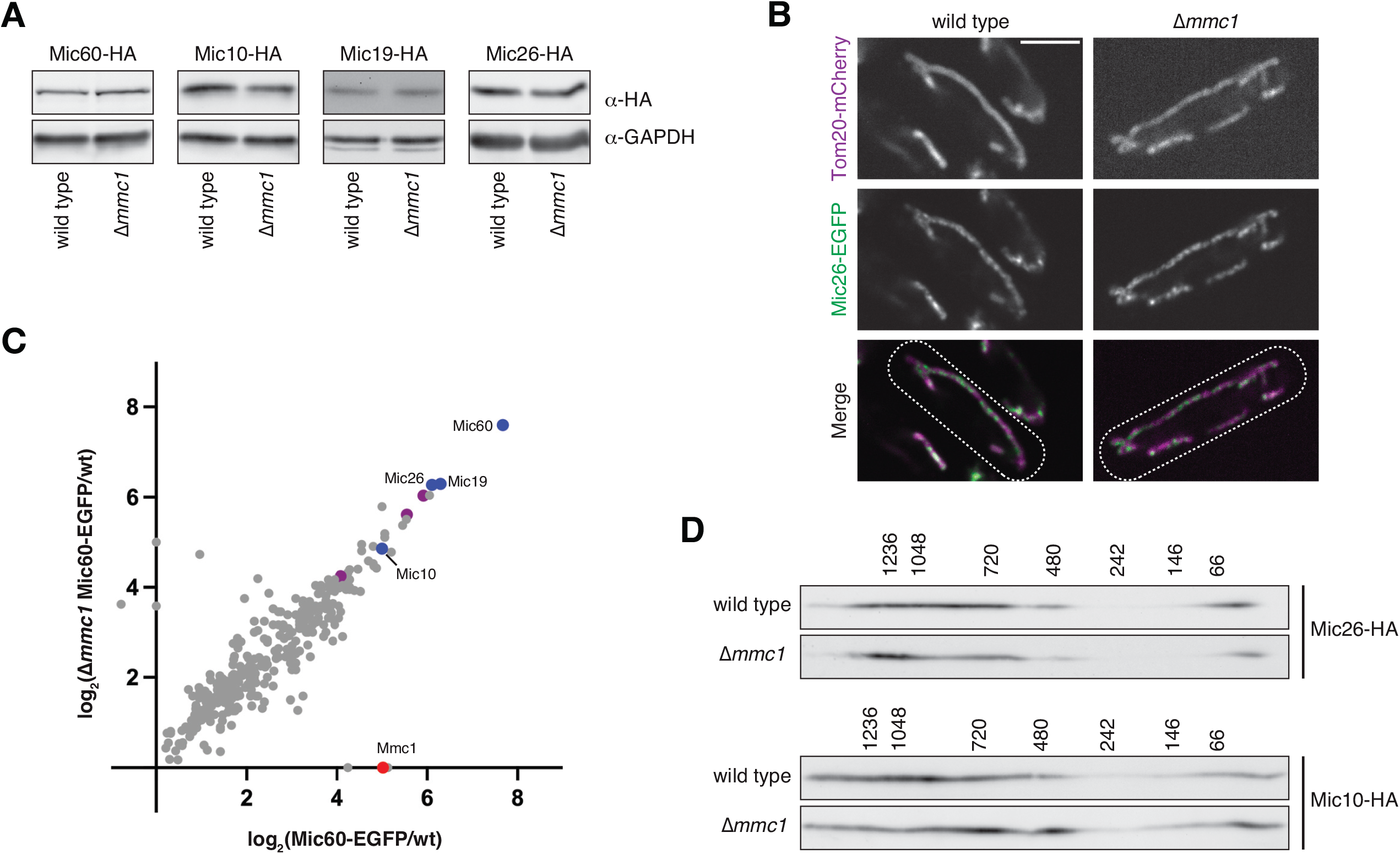
Mmc1 is not required for assembly of the MICOS complex. **(A)** Western analysis with the indicated antibodies of whole cell lysates from wild type or Δ*mmc1* cells expressing the indicated chromosomally HA-tagged MICOS subunit. **(B)** Single plane confocal microscopy images of wild type (left) or Δ*mmc1* (right) cells co-expressing chromosomally tagged Tom20-mCherry (magenta) and Mic26-EGFP (green) and grown to exponential phase in YES media. **(C)** A plot of the relative enrichment of total spectral matches of proteins identified from IP/MS of strains expressing Mic60-EGFP relative to a non-EGFP expressing strain in wild type cells (x-axis) or Δ*mmc1* cells (y-axis). Identified core MICOS subunits are highlighted in blue, MIB subunits are highlighted in magenta, and Mmc1 is highlighted in red. Data represent values from two independent experiments. See also Table S6. **(D)** Two-dimensional Blue Native PAGE (2D BN-PAGE) and Western analysis of mitochondria isolated from the indicated strains expressing chromosomally tagged Mic26-HA (top) or Mic10-HA (bottom). The molecular weight of assemblies as determined by the first dimension of BN-PAGE is displayed above the blots. Scale bar = 4 µm (B). Cell boundaries are indicated with dotted lines. See also Figure S5.

Next, we tested whether MICOS localization was affected in the absence of Mmc1. We imaged wild type and Δ*mmc1* cells co-expressing Mic26-EGFP and Tom20-mCherry by confocal microscopy. However, in the absence of Mmc1, Mic26-EGFP retained its punctate appearance, suggesting its distribution within mitochondria does not depend on Mmc1 (Fig. 5B). We also performed proteomic analysis of Mic60-EGFP in wild type and Δ*mmc1* cells, finding that Mic60 robustly retained its association with all MICOS and MIB subunits in the absence of Mmc1 (Fig. 5C, Supplemental Table 6). We then assayed whether loss of Mmc1 affected MICOS assembly by 2-Dimensional Blue Native PAGE (2D BN-PAGE). The tagged MICOS subunits Mic10-HA and Mic26-HA could be detected in large complexes ranging from ∼500 kDa to over ∼1.2 MDa in mitochondria isolated from fission yeast (Fig. 5D). Consistent with the proteomic analysis of Mic60-EGFP, the native sizes of Mic10 and Mic26 were unaffected in Δ*mmc1* cells (Fig. 5D). Together, these data suggest that the mitochondrial morphology defect observed in the absence of Mmc1 is not due to MICOS destabilization or assembly defects.

### Mmc1 and MICOS additively contribute to mitochondrial morphology

As Mmc1 does not appear to be required for MICOS assembly, yet co-localizes and associates with the MICOS complex, we considered that Mmc1 may work in concert with MICOS to generate cristae architecture. We therefore asked how combined loss of MICOS subunits and Mmc1 affected mitochondrial morphology. We generated Δ*mic10 Δmmc1* cells and Δ*mic60 Δmmc1* cells that co-expressed Tom20-EGFP and examined their mitochondria by confocal microscopy. Unlike the loss of individual MICOS subunits, which primarily led to lamellar mitochondrial morphology, the combined loss of Mmc1 and MICOS subunits caused a drastic increase in the presence of ring structures, including those that were over three microns in diameter (Fig. 6A-6B). In electron microscopy analysis, such structures could also be observed in Δ*mic60 Δmmc1* cells as enlarged versions of the less electron-dense internal structures found occasionally in Δ*mmc1* or Δ*mic60* cells (Fig. 5C).

**Figure 6.**
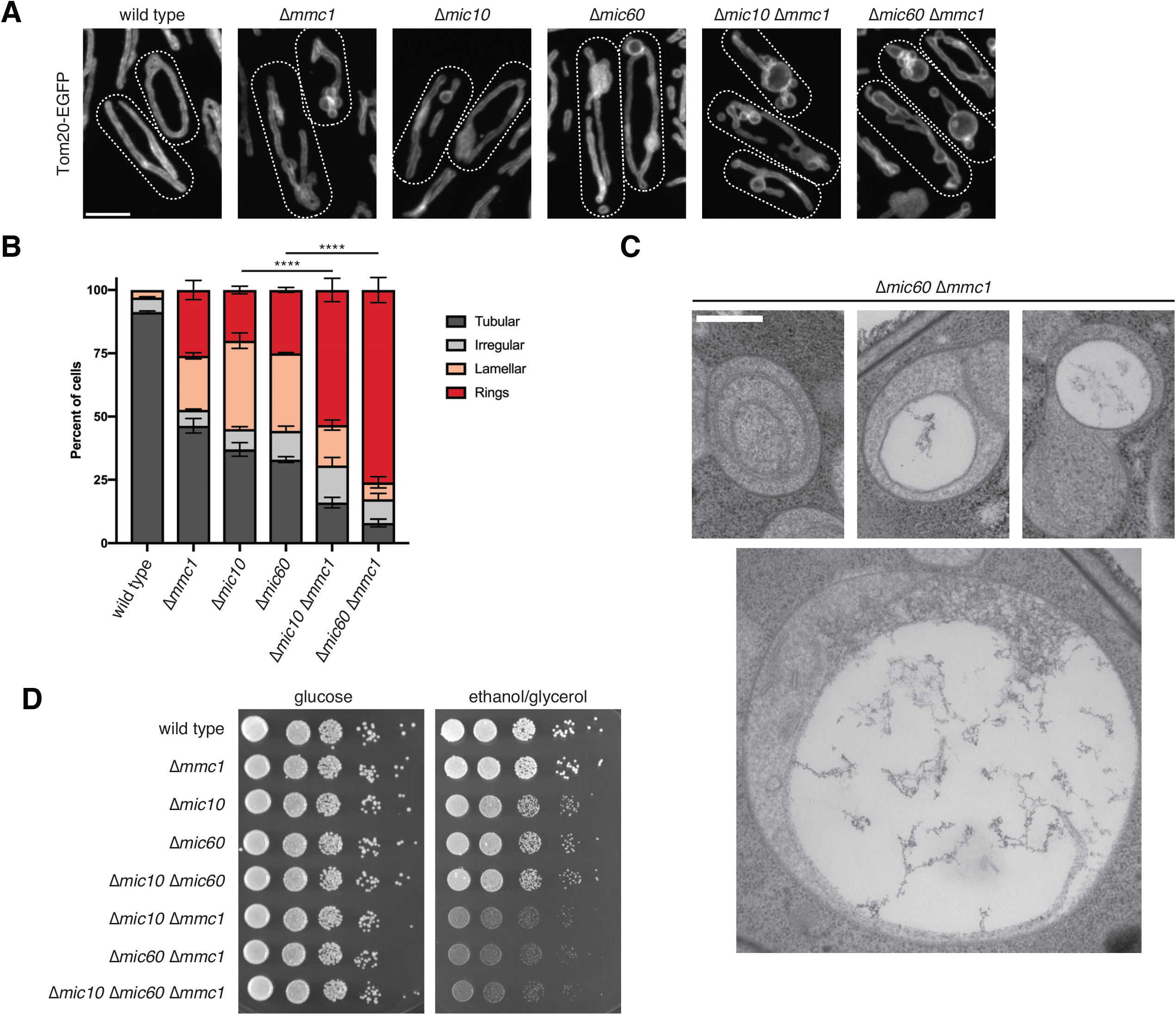
Mmc1 and MICOS additively contribute to mitochondrial morphology. **(A)** Maximum intensity projections of confocal images of the indicated strains expressing chromosomally tagged Tom20-EGFP and grown to exponential phase in EMM media. **(B)** A graph of the categorization of mitochondrial morphology from cells as in (A). Data shown represent 100 cells per strain in each of three independent experiments, and bars indicate S.E.M. Asterisks (****p<0.0001) represent two-way ANOVA with Tukey’s multiple comparisons test of tubular morphology. **(C)** Representative electron micrographs from Δ*mic60 Δmmc1* cells grown to exponential phase in EMM media. **(D)** Serial dilutions of the indicated strains grown to exponential phase in YES glucose media and plated on YES media containing glucose or the non-fermentable carbon source ethanol/glycerol and allowed to grow at 30°C for 3 days (glucose) or 9 days (ethanol/glycerol). Scale bars = 4 µm (A), 400 nm (C). Cell boundaries are indicated with dotted lines.

Next, we examined how the combined morphological defect of the loss of Mmc1 and MICOS subunits related to cellular respiratory growth. Strikingly, we observed a severe synthetic growth defect specifically on respiration-requiring media in the absence of Mmc1 and MICOS subunits (Fig. 6D). Importantly, combined loss of Mic10 and Mic60 did not additively affect mitochondrial respiratory growth either in the presence or absence of Mmc1 (Fig. 6D). These genetic interaction data indicate that while Mmc1 associates with the MICOS complex, it influences cristae morphology through a genetically parallel function.

### Mmc1 is a Dynamin Related Protein-like pseudoenzyme

To gain insight into the mechanism of Mmc1 contribution to cristae morphology, we next examined the sequence diversity of Mmc1 homologs. While there is no obvious Mmc1 ortholog in humans or *S. cerevisiae*, orthologs can be found widely throughout other fungal species (Fig. 7A, Fig. S6, Supplemental Table 7). Importantly, these Mmc1 orthologs, while distantly related at the primary sequence level, contain predicted mitochondrial targeting sequences (MTSs) and C-terminal transmembrane domains, suggesting they likely have similar localization and function as Mmc1.

**Figure 7.**
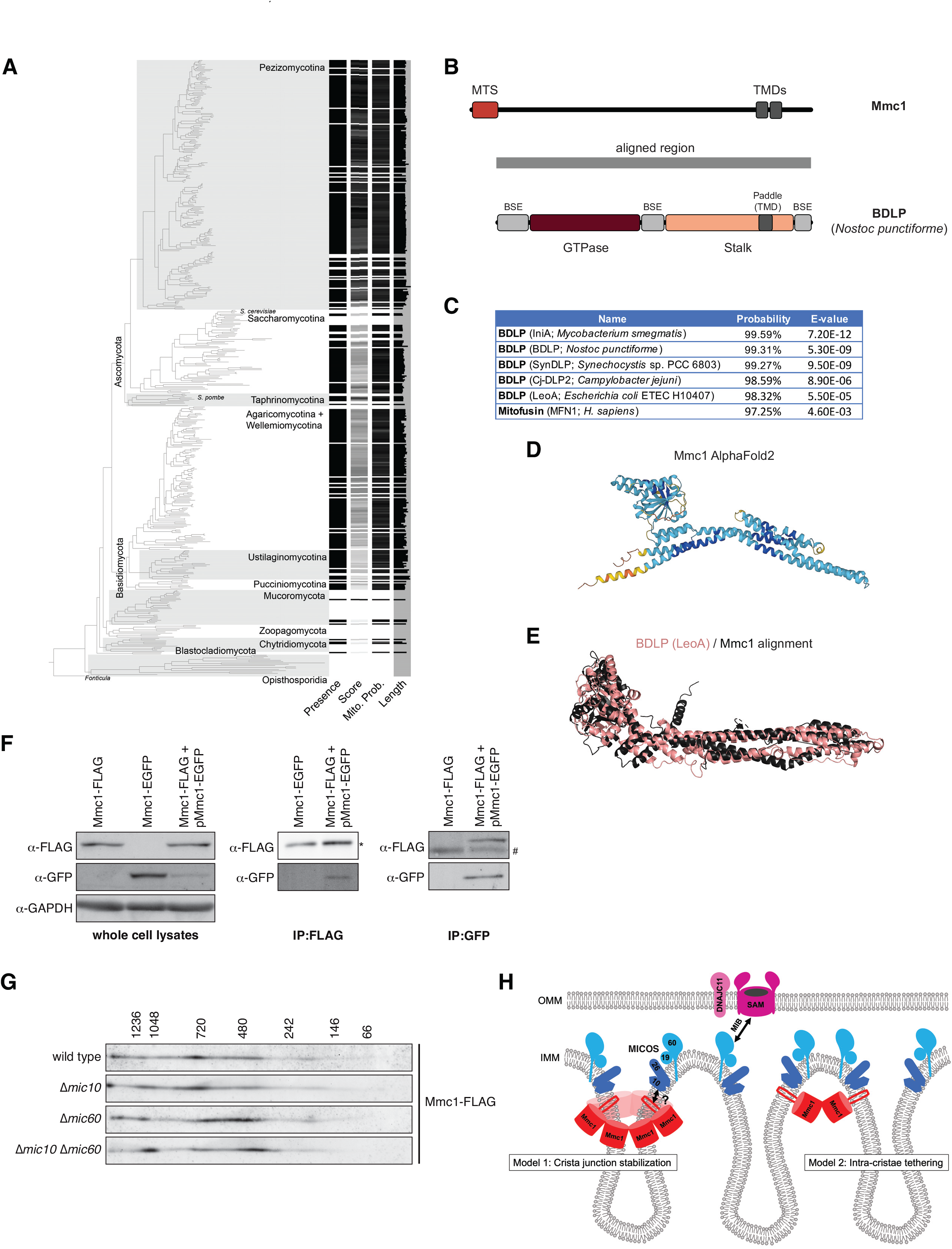
Mmc1 is a DRP-like pseudoenzyme and self-associates in MICOS-independent high molecular weight complexes. **(A)** Phylogenetic distribution of candidate Mmc1 homologs in the Fungi. Proteins were considered candidate orthologs if they were predicted to target to mitochondria and contain two C-terminal transmembrane segments. Score represents a measure of sequence similarity to a profile Hidden Markov Model built from iterative jackhammer searches using SpMmc1 as query against Mycocosm proteomes. Probability of mitochondrial targeting (Mito. Prob.) is indicated by grayscale shading with higher scores displayed in black. The relative lengths of identified homologs are displayed in horizontal bars (maximum length = 940 amino acids). For source data, see Table S7. For a detailed phylogenetic tree with species names see Figure S6. **(B)** A schematic is shown depicting the sequence alignment between Mmc1 and BDLP from *Nostoc punctiforme* as determined by HHPRED. The BDLP domain architecture is reproduced from Jimah et al^23^. See also File S1. **(C)** A table is shown of HHPRED analysis querying the Mmc1 sequence against the Protein Data Bank (PDB) database. HHPRED analysis was performed with global realignment. **(D)** The predicted AlphaFold2 structure of Mmc1. **(E)** Alignment of the Mmc1 AlphaFold2 structure (black) with the structure of (F) BDLP LeoA^49^ (PDB 4AUR_A) (pink). Alignment was performed with FATCAT^48^ with flexible modeling. **(F)** Western blot analysis is shown of whole cell lysates (left panel) and IP eluates from the indicated strains expressing chromosomally tagged Mmc1-FLAG or Mmc1-EGFP, or a strain co-expressing Mmc1-FLAG and plasmid-borne pMmc1-EGFP. IPs were performed with the indicated antibodies. The asterisk marks a band consistent with the size of IgG heavy chain, which migrates at a similar size to Mmc1-FLAG. The hashtag marks a non-specific band. **(G)** 2D BN-PAGE and Western analysis of mitochondria isolated from the indicated strains expressing chromosomally tagged Mmc1-FLAG. **(H)** A model depicting potential roles for Mmc1 in promoting cristae maintenance. See also Figure S7.

To explore a potential molecular function for Mmc1, we performed an *in silico* analysis. Using InterPro/Pfam^43^ to identify protein family membership, no sequence-conserved functional domains were identified within Mmc1. We then used HHPRED^45^ to identify structurally similar proteins that may be remotely related to Mmc1. Notably, when Mmc1 was queried against the Protein Data Bank (PDB) of known protein structures, several members of the Dynamin Related Protein (DRP) family were significant hits (Fig. 7B-7C). DRPs are large GTPases that hydrolyze GTP to perform membrane remodeling processes, including mitochondrial fission and fusion dynamics^23,24^. The top hits for Mmc1 were Bacterial Dynamin-Like Proteins (BDLPs), including *Mycobacterium smegmatis* IniA and *Nostoc punctiforme* BDLP. Notably, the human mitochondrial DRP mitofusin 1 (MFN1) was also identified as a significant hit (Fig. 7C). Importantly, the Mmc1 sequence showed similarity with each DRP hit across the entire mature protein sequence, including the GTPase and stalk domains (Fig. 7B; Supplemental File 1).

In addition to distant sequence-based homology, the AlphaFold2^46,47^ predicted structure of Mmc1 appears similar to that of solved DRP structures, including an N-terminal fold that resembles a GTPase domain and C-terminal alpha-helical regions similar to the helical stalk region of DRPs (Fig. 7D). The flexible structural alignment program FATCAT^48^ also detected significant similarity between Mmc1 and BDLPs and was able to align the AlphaFold2 structure of Mmc1 with the solved structures of LeoA^49^ (Fig. 7E) and IniA^50^ (Fig. S7A). Further, despite their sequence divergence relative to Mmc1, each ortholog we examined has predicted DRP-like folds by both HHPRED and AlphaFold2 analyses (Fig. S7B-S7C). Importantly, however, Mmc1 is a pseudoenzyme as the critical sequence motifs required for GTP binding and hydrolysis present in all DRP family members are absent in Mmc1 and its orthologs (Fig. S7D). Thus, while Mmc1 has distant homology and is predicted to share structural similarity with DRPs, the protein likely does not bind or hydrolyze GTP to remodel membranes. The loss of GTPase activity also may help explain the weak sequence conservation among apparent orthologs, as there is no longer evolutionary pressure to maintain the active site.

### Mmc1 self-associates in high molecular weight complexes

A common feature of DRPs is their ability to self-associate in oligomeric structures that allow them to perform their functions in membrane remodeling^23^. To assess whether Mmc1 shared characteristics with DRP family members, we first tested whether Mmc1 was capable of interacting with itself. To address this, we expressed plasmid-borne Mmc1-EGFP in a cell expressing chromosomally tagged Mmc1-FLAG, immunoprecipitated with either anti-FLAG or anti-GFP, and performed Western analysis. In both cases, Mmc1-FLAG and Mmc1-EGFP were able to reciprocally interact (Fig. 7F). These data suggest that Mmc1 can self-associate in cells, though this interaction may be mediated through indirect interactions with MICOS subunits.

Next, we wanted to ascertain whether Mmc1 assembles into high molecular weight structures, similar to DRP family members. We performed 2D BN-PAGE analysis of mitochondria purified from Mmc1-FLAG expressing cells. As in the case of MICOS subunits, Mmc1 (monomer: ∼60 kD) assembled in large complexes ranging in size from ∼400 kDa to over 1.2 MDa (Fig. 7G). Notably, the size of Mmc1 assemblies was largely unchanged in the absence of either Mic10, Mic60, or in *Δmic10 Δmic60* cells where the MICOS complex is likely completely destabilized (Fig. 7G and Fig. S2). Together, these data suggest that like DRP family members, Mmc1 self-associates in high molecular weight complexes, and does so independent of the MICOS complex.

## Discussion

While the precise mechanisms by which MICOS contributes to cristae architecture are not completely understood, the complex is thought to work locally at cristae junctions. Here we show that in the fission yeast *Schizosaccharomyces pombe*, MICOS cooperatively contributes to cristae architecture with the matrix-facing IMM protein Mmc1. We find that Mmc1 interacts with MICOS subunits and concentrates in proximity to MICOS at cristae junctions. However, Mmc1 is not required for MICOS stability or assembly and instead works additively with MICOS to promote cristae morphology. Mmc1 is phylogenetically widespread among fungi but has sporadically been lost in a few species such as *Saccharomyces cerevisiae*. Thus, our data suggest that Mmc1 represents a new class of proteins that is ancestral to most of the Fungi and that work with MICOS to modulate mitochondrial cristae morphology.

Remote homology searches, AlphaFold2 structural predictions, and detailed sequence analysis indicate that Mmc1 resembles numerous members of the DRP family. Several DRP proteins are involved in mitochondrial dynamics and shaping, including the OMM-localized fusion and fission machines, mitofusins (Fzo1 in budding yeast) and Drp1 (Dnm1 in budding yeast), respectively^51^. Additionally, IMM fusion depends on the IMS-facing DRP family member OPA1 (Mgm1 in budding yeast). A common feature of these and other DRP proteins is that they self-assemble and remodel membranes in a GTP hydrolysis-dependent manner. Our data indicate that, like DRPs, Mmc1 can self-associate and form high molecular weight complexes. However, Mmc1 and its orthologs lack residues required for GTP binding and hydrolysis, suggesting it is part of an emerging class of GTPase pseudoenzymes^52^. Thus, Mmc1 may fold in a similar shape to DRPs, though likely does not generate force through a conformational change driven by GTPase activity. A potential model to explain a role for Mmc1 is that it utilizes its putative DRP-like fold to promote stability of membrane curvature, potentially by acting locally on the matrix side of the cristae junction (Fig. 7H). An alternative model is that Mmc1 may influence cristae architecture by mediating the relative positioning of cristae by *cis* interactions on adjacent cristae (Fig. 7H). The latter may potentially explain our observations in human cells that exogenously expressed Mmc1 causes the redistribution of MICOS complexes (Fig. 4).

While our data indicate that Mmc1 associates with and localizes in proximity to the MICOS complex, it remains to be determined the mechanism and function of this association. Mmc1 can form high molecular weight assemblies independently of MICOS and, based on their additive effects, Mmc1 can contribute to cristae morphology independently. However, we observe that Mmc1 co-localizes with the Mic10/Mic26 subcomplex, which forms foci in the absence of Mic60 or Mic19. Additionally, we find that loss of Mic10 or Mic26 leads to disruption of Mmc1 focal structures. These data suggest that the physical proximity of Mmc1 and MICOS is due to physical interaction, likely via the Mic10/Mic26 subcomplex, rather than through independent cues. Notably, Mmc1 is negligibly exposed to the IMS and likewise, Mic10/Mic26 are minimally exposed to the matrix. One possibility is that transmembrane domain associations between the proteins maintain their association (Fig. 7H), and a focus of future work will be to identify mutations that selectively disrupt the MICOS-Mmc1 interaction.

An outstanding question is whether other proteins perform a functionally conserved role in metazoans and other fungal species where Mmc1 is not found. Interestingly, a recent preprint identified a DRP family member in trypanosomes (*Tb*DBF*/Tb*MFNL) and several other metazoans with a similar domain organization to Mmc1, including an N-terminal MTS and C-terminal transmembrane domains^53^. Unlike Mmc1, *Tb*DBF has a canonical GTPase domain, and the protein has been implicated in fission/fusion dynamics^53,54^. However, given its topology, it will be interesting to determine if *Tb*DBF may additionally or instead work with MICOS to contribute to cristae architecture. In human cells where no such matrix-localized DRP has been identified, such a cristae-stabilizing scaffold could conceivably be performed by other structurally distinct factors. For example, recent work identified a BAR-domain containing protein that localizes to the mitochondrial matrix and is required for mitochondrial ultrastructure^55^. An alternative potential explanation is that the presence of Mmc1 is a feature of organisms, like *S. pombe*, that express fewer core MICOS subunits. It will be interesting to explore the possibility that the role of Mmc1 can be performed by MICOS complexes that have expanded to include additional subunits, for example, the presence of Mic12 homologs.

## Supplemental Data

**Figure S1.**
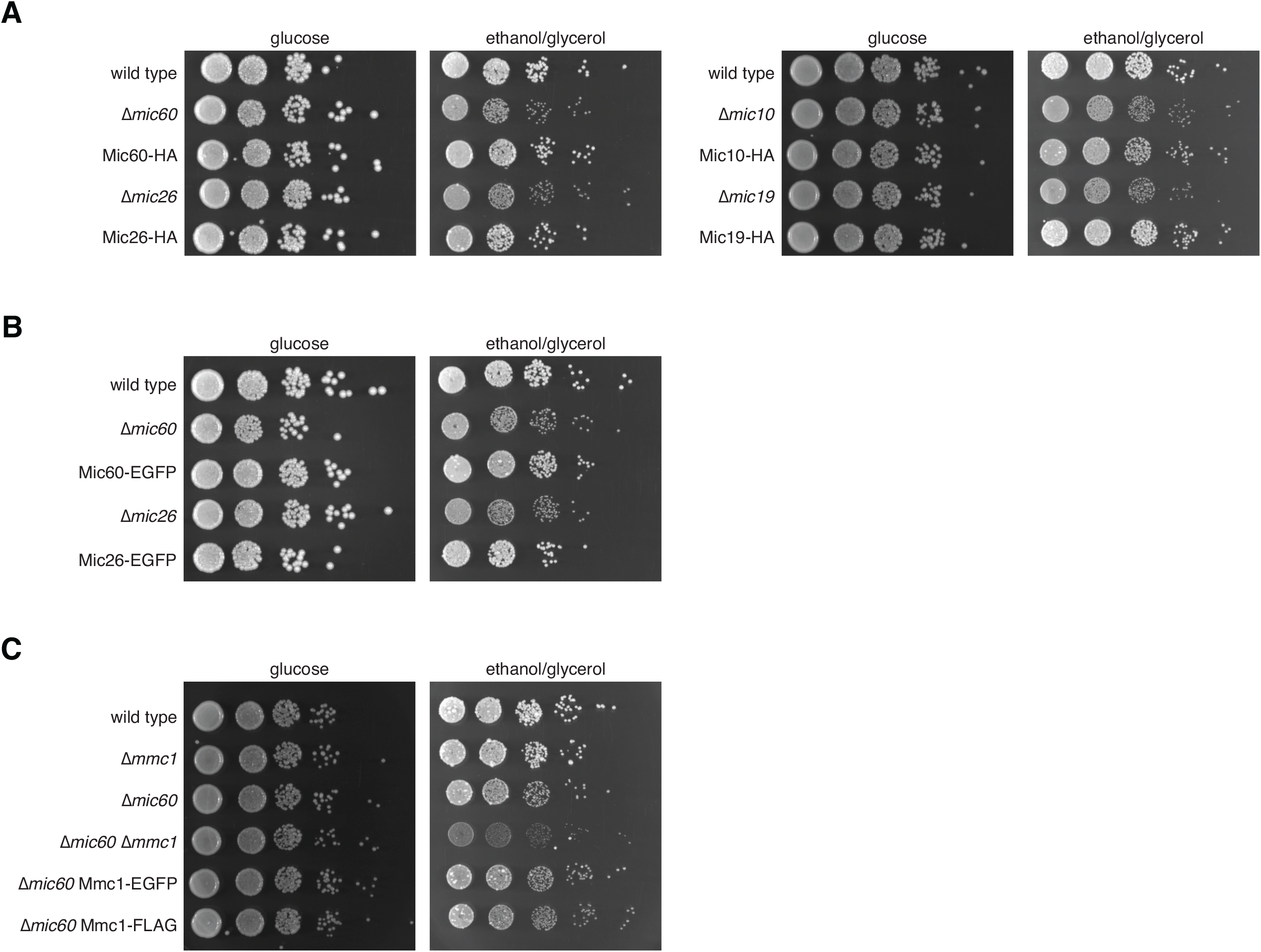
Tagged *S. pombe* MICOS subunits and Mmc1 are functional to maintain respiratory growth. **(A-B)** Serial dilutions of the indicated deletion strains or strains expressing the indicated chromosomally tagged MICOS subunits grown to exponential phase in YES glucose media and plated on YES media containing glucose or the non-fermentable carbon source ethanol/glycerol and allowed to grow at 30°C for 2-3 days (glucose) or 8-10 days (ethanol/glycerol). **(C)** As in (A-B) for cells expressing chromosomally tagged Mmc1-EGFP or Mmc1-FLAG. Related to Figures 1 and 2.

**Figure S2.**
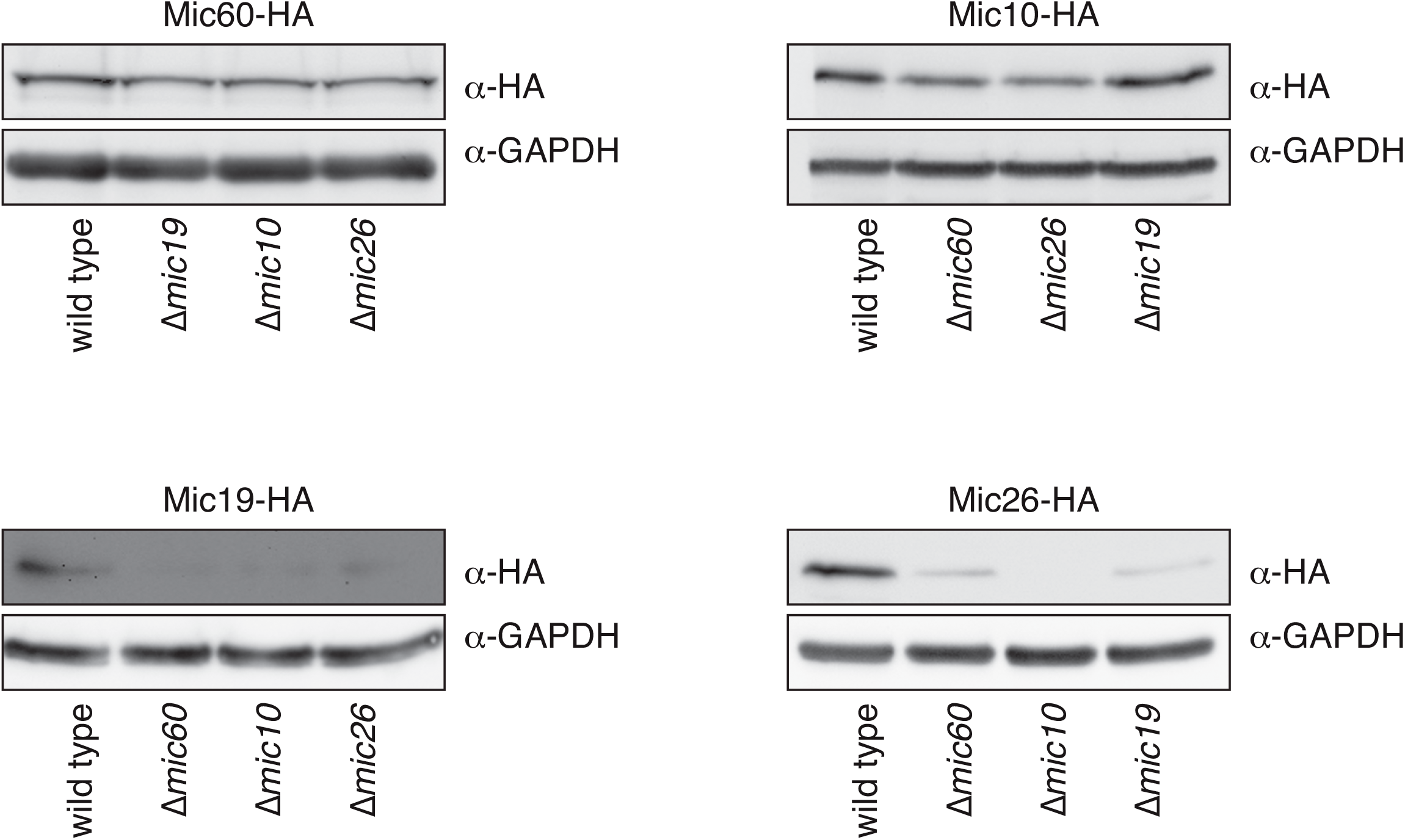
*S. pombe* MICOS subunits exhibit protein stability interdependence. Western analysis is shown with the indicated antibodies of whole cell lysates from the indicated strains expressing the indicated chromosomally HA-tagged MICOS subunit. Related to Figure 1.

**Figure S3.**
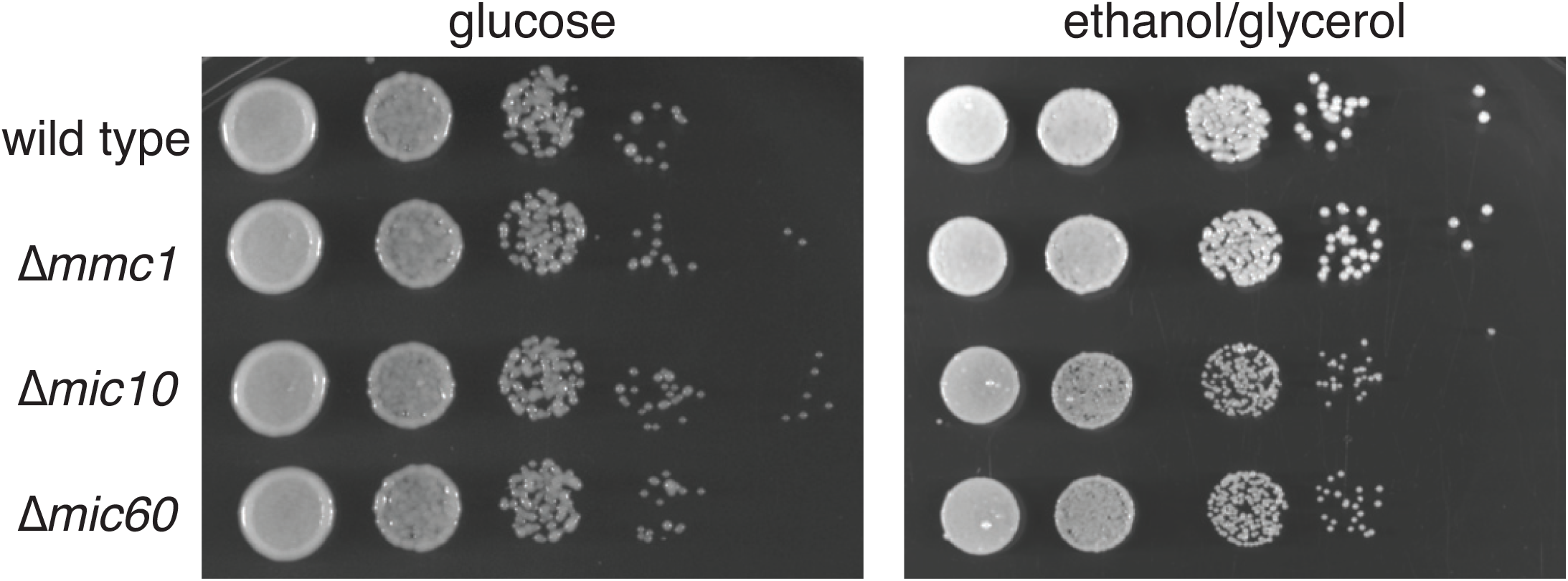
Loss of Mmc1 does not grossly affect the rate of cell growth on respiration-requiring media. Serial dilutions of the indicated deletion strains grown to exponential phase in YES glucose media and plated on YES media containing glucose or the non-fermentable carbon source ethanol/glycerol and allowed to grow at 30°C for 2 days (glucose) or 9 days (ethanol/glycerol). Related to Figure 2.

**Figure S4.**
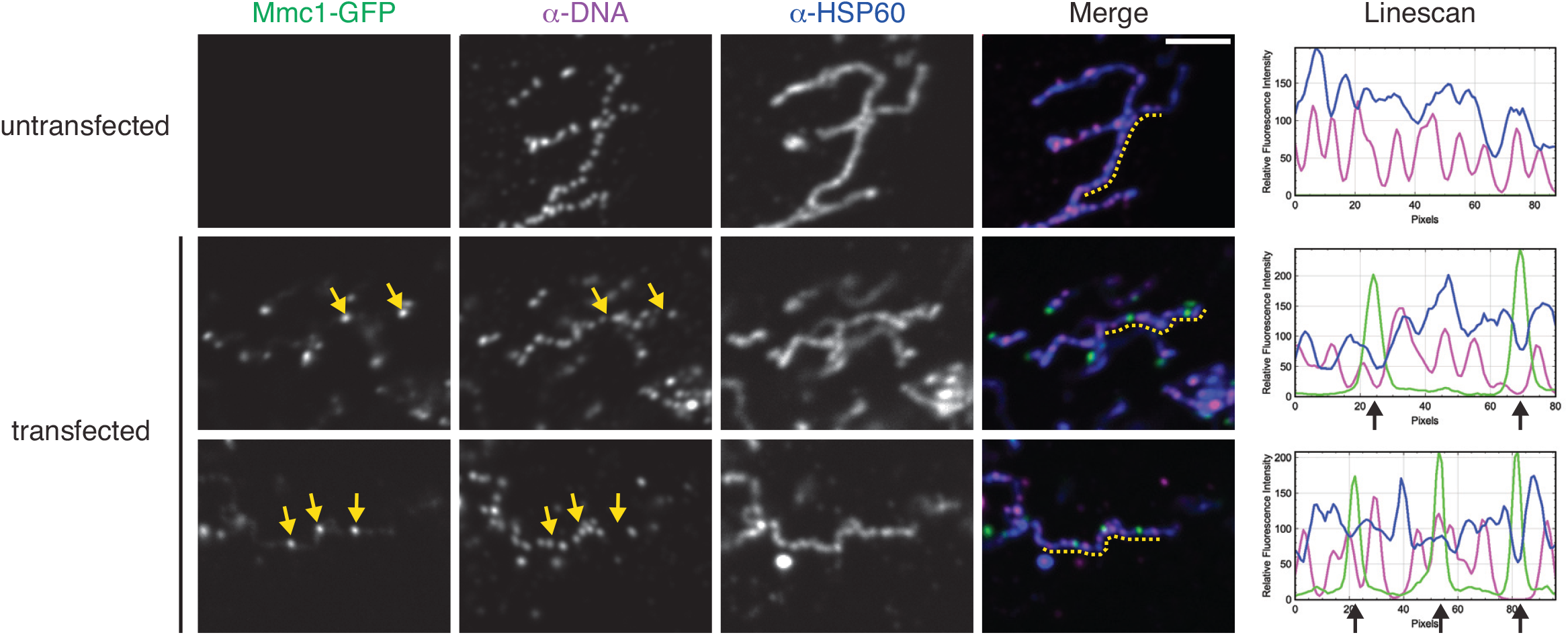
Mmc1 focal structures in human cells do not co-enrich with nucleoids. Single plane confocal fluorescence microscopy images are shown of a non-transfected U2OS cell (top) or cells transfected with Mmc1-GFP (green). Cells were fixed and immunolabeled with anti-dsDNA (magenta) and anti-HSP60 (blue). Fluorescence intensity linescans shown at right were performed corresponding to the dashed yellow line shown on the merge image. Yellow arrows mark Mmc1 focal enrichments that correspond to the indicated location on the mtDNA image and the associated black arrows that are marked on linescans. Related to Figure 4.

**Figure S5.**
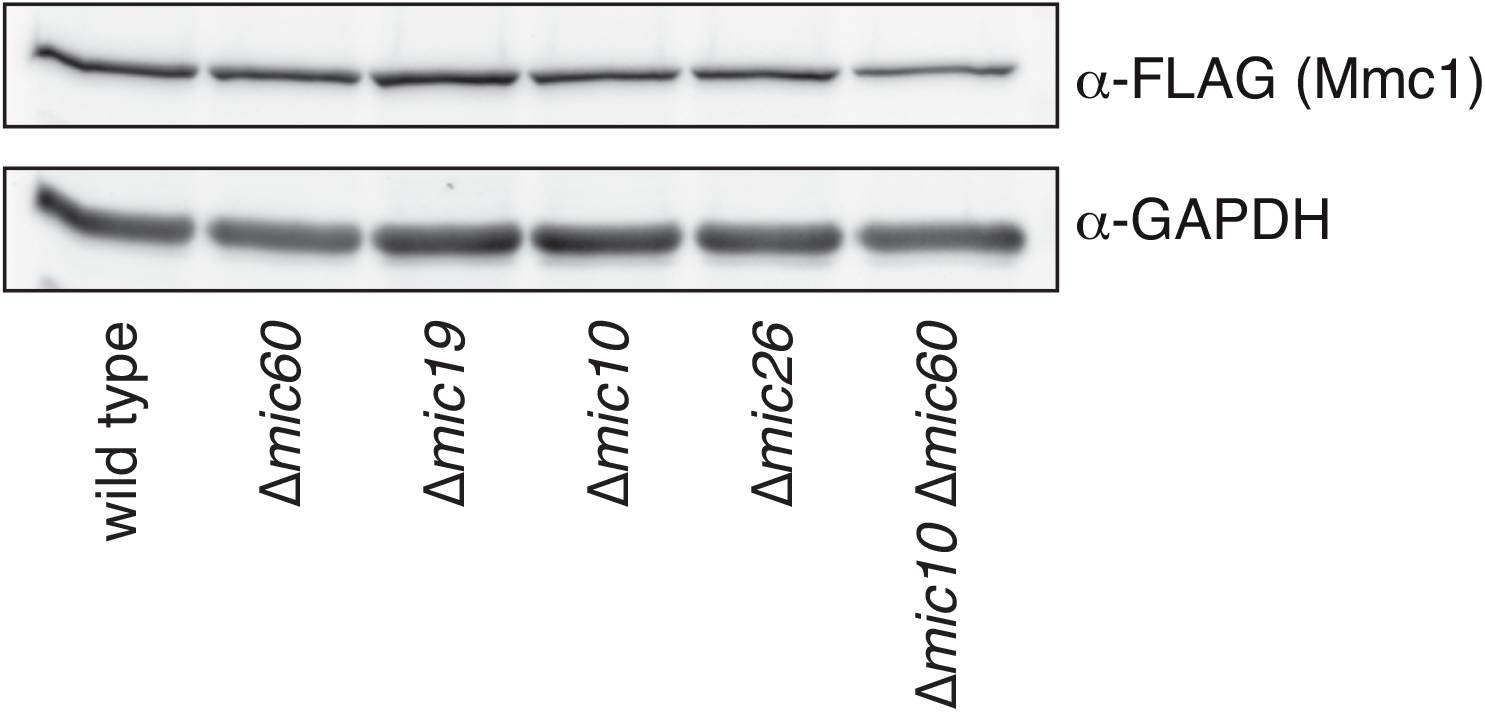
Mmc1 protein expression levels are not grossly affected by the loss of MICOS subunit. Western analysis is shown with the indicated antibodies of whole cell lysates from the indicated strains expressing the chromosomally tagged Mmc1-FLAG. Related to Figure 5.

**Figure S6.**
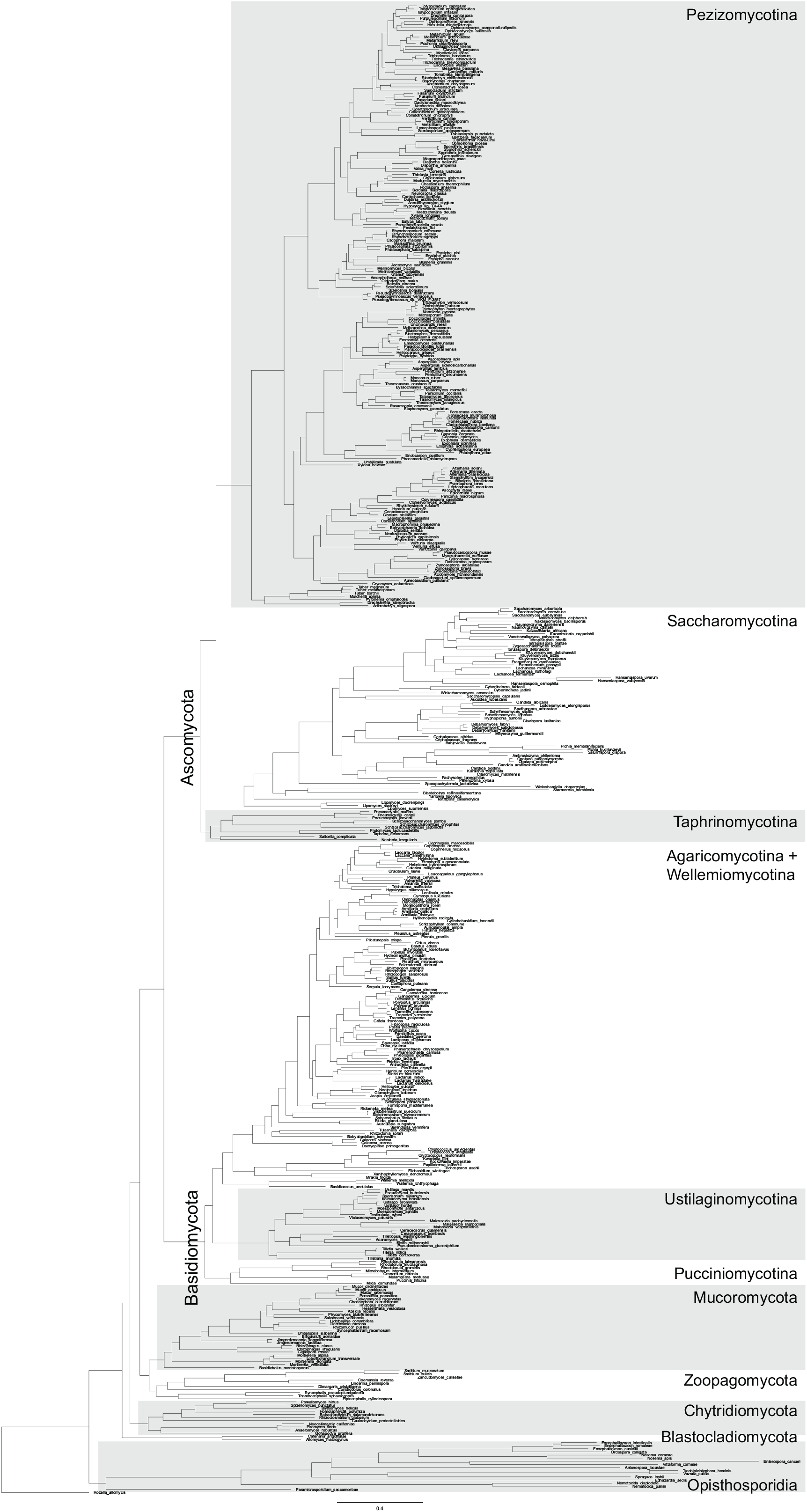
A labeled phylogenetic tree of Fungi used to display the phylogenetic distribution of putative Mmc1 orthologs. A phylogenetic tree with complete species names of fungi investigated corresponding to Figure 7A and Table S7.

**Figure S7.**
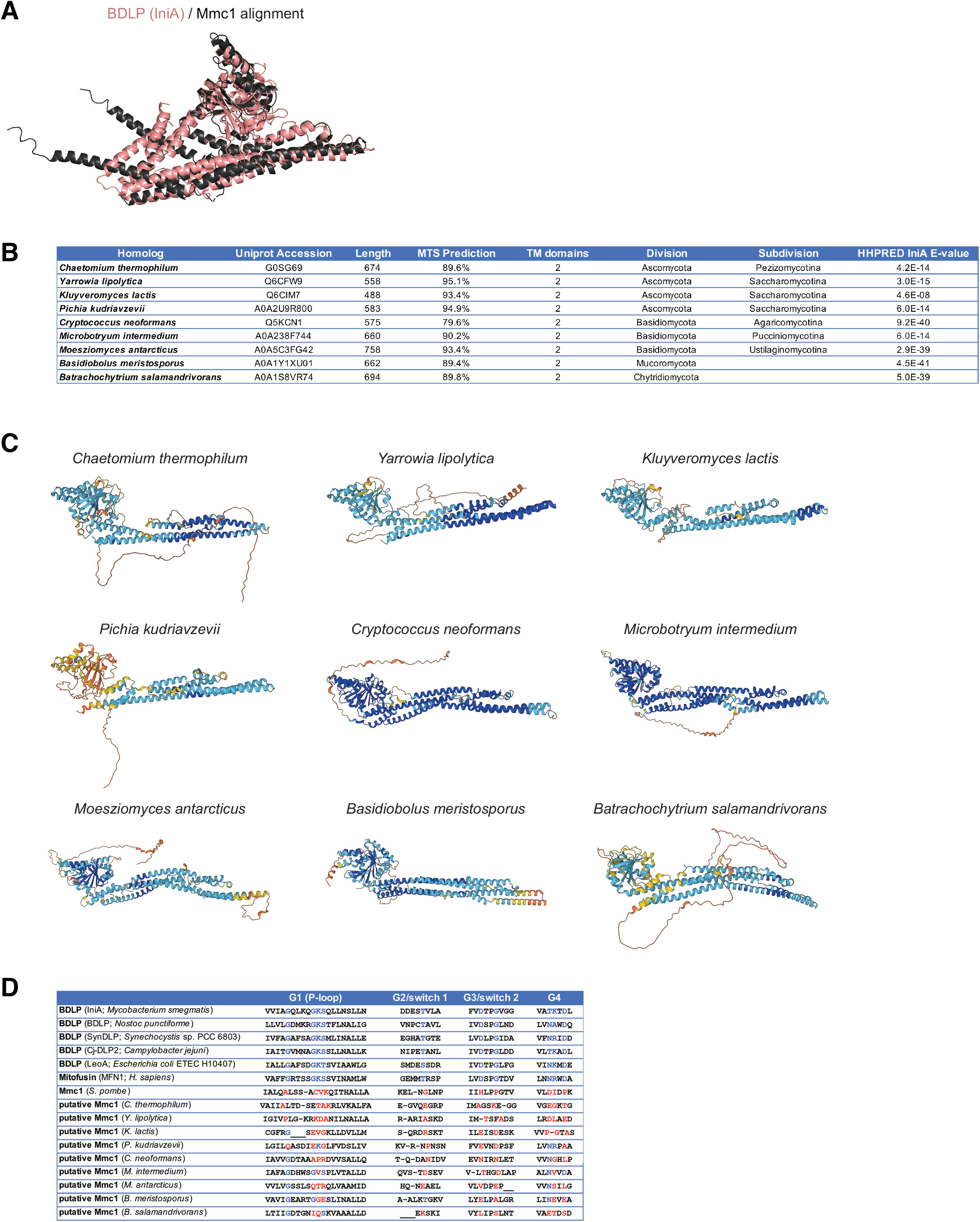
Mmc1 orthologs are widely found in fungal species but do not have key residues required for GTP binding or hydrolysis. **(A)** An alignment is shown of the Mmc1 AlphaFold2 structure (black) with the structure of the BDLP IniA^50^ (PDB 6J72_A) (pink). The alignment was generated using FATCAT^48^ with flexible modeling. **(B)** A table is shown of sample Mmc1 orthologs identified in the phylogenetic analysis. The protein sequence was used to query the Protein Data Bank using HHPRED and the E-value for homology to the BDLP IniA is shown. **(C)** The AlphaFold2 predicted structure is shown for each Mmc1 ortholog listed in (B). Structures were oriented so that the GTPase-like fold of each ortholog faces the top-left of the image. **(D)** A table is shown depicting conserved residues (blue) required for GTP binding and hydrolysis among hit DRP proteins identified in HHPRED analysis. The corresponding residues of Mmc1 and putative homologs from (B) are shown based on the HHPRED alignment of each to the group of DRPs. Residues are shown in red when they differ from that of consensus GTPases. Dashes indicate sequence omitted from the alignment and underscores mark gaps in sequence. Related to Figure 7.

**File S1. HHPRED alignment of Mmc1 and DRP family members.** A multiple sequence alignment file of Mmc1 and DRP family members identified by HHPRED when queried against the PDB with global realignment.

## Materials and Methods

### S. pombe growth and strain construction

Wild type haploid *S. pombe* was a generous gift from Mike Henne (UT Southwestern). *S. pombe* strains used in this study were routinely cultured in YES (0.5% yeast extract, 3% glucose, 225 mg/L adenine, 225 mg/L leucine, 225 mg/L histidine, 225 mg/L uracil, 225 mg/L lysine). Where indicated, cells were grown in EMM media for imaging (Sunrise Science; supplemented with 225 mg/L adenine, 225 mg/L leucine, 225 mg/L histidine, 225 mg/L uracil, 225 mg/L lysine, 20 mg/L thiamine). For respiratory growth, cells were plated on solid YES ethanol/glycerol media (0.5% yeast extract, 3% glycerol, 3% ethanol, 225 mg/L adenine, 225 mg/L leucine, 225 mg/L histidine, 225 mg/L uracil, 225 mg/L lysine). All strains used in this study were stored at −80°C and freshly revived immediately prior to experiments.

Strains were constructed by PCR-based homologous recombination and lithium acetate transformation as described^56^, except cells were incubated for 5 h in growth media after heat shock treatment for recovery before plating on selection media. Deletions were generated by replacing the entire open reading frame with the NatMX6, HphMX6, or KanMX6 cassettes from pFA6a-series plasmids following lithium acetate transformation^56^. C-terminal tagging of proteins at the indicated endogenous chromosomal locus was accomplished using pFA6a-link-yEGFP- Kan, pFA6a-mCherry-HphMX6, pFA6a-mCherry-NatMX, pFA6a-yoHalo-HphMX6, pFA6A- 3xFLAG-Kan, or pFA6a-3xHA-Kan^25,44,57,58^. pFA6a-3xFLAG-Kan was generated by cloning the Kan cassette into the BglII/EcoRV sites of pFA6a-3xFLAG-His^59^, replacing the His cassette. pFA6a-yoHalo-HphMX6 was generated by cloning the HphMX6 cassette into the BglII/EcoRV sites of pFA6a-yoHalo::CaUra3^60^.

An Mmc1-EGFP plasmid (pMmc1-EGFP) for expression in *S. pombe* was generated by PCR amplifying: (1) the Mmc1 coding sequence with its promoter region, (2) the yEGFP coding sequence, and (3) the ADH terminator sequence, and cloning into mitoRED::Hyg^61^ by Gibson assembly, replacing the mito-mCherry cassette. The pMmc1-GFP expression plasmid was linearized and integrated into the *S. pombe* genome at the *leu1* locus.

### Confocal fluorescence microscopy and analysis

All confocal fluorescence microscopy was carried out on a Nikon Spinning Disk Confocal microscope with Yokogawa CSU-W1 SoRa and equipped with a Hamamatsu Orca-Fusion sCMOS camera, a Nikon 100x 1.45 NA objective, and an environmental chamber. Images were acquired with Nikon Elements software using the standard spinning disk module and all z-series images were obtained with a 0.2 μm step size. *S. pombe* images were acquired at 30°C. Images of U2OS cells were acquired at 37°C.

*S. pombe* strains were grown to exponential phase at 30°C in the growth media indicated in figure legends, harvested, and immobilized on a 3% agarose bed of EMM on cavity microscope slides for imaging. Cells expressing HaloTag were treated with JF646 HaloTag ligand (Promega; 1 μM final concentration in growth medium) for 30 minutes at room temperature, washed once with growth media, concentrated, and imaged. Where indicated, cells were labeled with 200 nM Mitotracker Deep Red FM (Thermo Fisher Scientific) for 30 min, washed three times with growth media, concentrated, and imaged. Multiple fields of view were imaged per strain or condition in at least three independent experiments, and ImageJ/Fiji was used to make linear adjustments to representative images. Linescan analysis was performed with the RGB Profiles Tool macro (https://imagej.nih.gov/ij/macros/tools/RGBProfilesTool.txt) with slight modifications.

To examine mitochondrial morphology or the presence of Mmc1 foci, strain identity was blinded before assessment, and cells were manually categorized as indicated in figures and legends. In the case of cells with mixed mitochondrial morphologies, cells were categorized with the more prominent morphology.

### Analysis of S. pombe mitochondrial ultrastructure

For electron microscopy analysis, *S. pombe* strains were grown to exponential phase at 30°C in EMM growth media. Cultures were harvested and prepared for electron microscopy as described previously^62^. After high-pressure freezing in a Wohlwend Compact 03 high-pressure freezer, the samples were freeze substituted in 2% OsO_4_, 1% glutaraldehyde, 0.1% uranyl acetate, and 5% water in acetone using a Leica AFS2 freeze substitution apparatus. After freeze substitution, the samples were warmed to room temperature, dehydrated through 100% acetone, and embedded in Spurr’s resin. Thin sections were cut using a Leica UC7 ultramicrotome, poststained with uranyl acetate and lead citrate, and imaged on a JEOL 1400 Flash transmission electron microscope.

### *S. pombe* growth analysis

Respiratory growth of *S. pombe* was assayed as described previously^63^. Strains were grown exponentially at 30°C in YES. Cells were harvested by centrifugation (6000 g for 2 min at room temperature) followed by resuspending in water at a density of 0.5 OD_600_/ml. A 10-fold serial dilution of the cells was spotted on YES media containing glucose or ethanol/glycerol as carbon sources (see *S. pombe* growth above). Plates were incubated at 30°C for the time duration indicated in figure legends.

### Immunopurifications and downstream analyses

Immunopurifications were performed as described^26^ except 500 OD_600_ of *S. pombe* cells that were grown to exponential phase in YES media were used. Cells were harvested and pellets were rinsed with water and resuspended in lysis buffer (20 mM HEPES pH 7.4, 150 mM KOAc, 2 mM Mg(Ac)_2_, 1 mM EGTA, 0.6 M sorbitol, and 1× protease inhibitor cocktail [539131; MilliporeSigma]), flash-frozen dropwise in liquid nitrogen, and lysed using a Freezer/Mill (SPEX). Cell lysates were cleared by centrifugation, cross-linked (except where indicated below) for 30 min with 1 mM DSP (Thermo Fisher Scientific), solubilized with 1% digitonin (EMD Millipore) for 30 min, and pelleted again. The resulting supernatant was used for purification with μMACS anti-GFP microbeads (Miltenyi Biotech), and beads were isolated with μ columns and a μMACS separator (Miltenyi Biotech). Samples were eluted using on-bead trypsin digestion by adding 25 μl Elution Buffer 1 (2 M urea, 50 mM Tris-Cl pH 7.5, 1 mM DTT, 5 μg/ml trypsin) for 30 min followed by two additions of 50 μl Elution Buffer 2 (2 M urea, 50 mM Tris-Cl pH 7.5, 5 mM chloroacetamide) and incubated overnight. 1 μl of trifluoroacetic acid was applied to quench the reaction and samples were submitted to the UT Southwestern Proteomics Core for liquid chromatography/tandem MS analysis.

For self-interaction of Mmc1, immunoprecipitations were carried out using the above protocol, except without the addition of a crosslinker. Proteins were eluted with 2 × 30 μl of Laemmli buffer pre-warmed to 95°C. 30% of the total elute was loaded on an SDS-PAGE gel and analyzed by Western blotting as described below.

For mass spectrometry analysis, the samples underwent solid-phase extraction cleanup with an Oasis HLB plate (Waters) and were subsequently dried and reconstituted into 10 µl of 2% ACN and 0.1% trifluoroacetic acid. 2 µl of these samples were injected onto a QExactive HF or an Orbitrap Elite mass spectrometer coupled to an Ultimate 3000 RSLC-Nano liquid chromatography system. Samples were injected onto a 75-µm-inner-diameter, 15-cm-long EasySpray column (Thermo Fisher Scientific) and eluted with a gradient from 0 to 28% buffer B over 90 min with a flow rate of 250 nl/min. Buffer A contained 2% (vol/vol) ACN and 0.1% formic acid in water, and buffer B contained 80% (vol/vol) ACN, 10% (vol/vol) trifluoroethanol, and 0.1% formic acid in water. The mass spectrometer operated in positive ion mode with a source voltage of 1.5-2.5 kV and an ion transfer tube temperature of 275°C. MS scans were acquired at 120,000 resolution in the Orbitrap, and ≤20 tandem MS spectra were obtained for each full spectrum acquired using higher-energy collisional dissociation for ions with charges 2–8. Dynamic exclusion was set for 20 s after an ion was selected for fragmentation.

Raw MS data files were analyzed using Proteome Discoverer v2.4 or v3.0 (Thermo Fisher Scientific), with peptide identification performed using Sequest HT searching against the *Schizosaccharomyces pombe* protein database from UniProt. Fragment and precursor tolerances of 10 ppm and 0.02 Da were specified, and three missed cleavages were allowed. Carbamidomethylation of Cys was set as a fixed modification, with oxidation of Met set as a variable modification. The false discovery rate cutoff was 1% for all peptides.

For all IP/MS analyses, each independent replicate was performed simultaneously with an untagged wild type strain. Except where described below, identified proteins were included in subsequent analysis if at least 15 total peptide spectral matches were identified in any individual replicate of either control or EGFP-tagged samples. For those proteins where no peptide spectral matches were identified, an imputed value of ‘1’ was given. The fold change of total spectral matches identified in the EGFP-tagged strain relative to the untagged strain was determined and the average fold change between experimental replicates is shown graphically in figures. The p-value was calculated by a two-tailed *t* test comparing the non-imputed EGFP-tagged spectral matches to the untagged spectral matches for each replicate.

For IP/MS analysis of Mmc1-EGFP in wild type versus Δ*mic10* and Δ*mic60* strains, two independent replicates were performed and compared with an untagged wild type strain. Identified proteins were included in subsequent analysis if (1) the protein was annotated as mitochondrial and (2) peptide spectral matches were enriched in the Mmc1-EGFP sample at least five-fold relative to the untagged wild type strain (with an imputed value of ‘1’ for unidentified proteins). Samples were normalized by the total peptide spectral matches identified, and for each analyzed protein, the fold change of total spectral matches identified by Mmc1-EGFP in Δ*mic10* or Δ*mic60* relative to wild type cells was determined.

### Mitochondrial isolation

Mitochondria were isolated as previously described with slight modifications^64^. Briefly, 500–1000 OD_600_ of *S. pombe* cells were grown to exponential phase in YES, harvested, and washed with water. Spheroplasts were made by digesting yeast cell walls with 4 mg/ml zymolyase 20T (Sunrise Science) in 1.2 M sorbitol. Spheroplasts were rinsed and resuspended in a minimal volume of cold MIB buffer (0.6 M sorbitol, 50 mM KCl, 20 mM HEPES pH 7.4) and homogenized with a glass dounce. Low-speed centrifugation (2400 g, 5 min, 4°C) was used to remove unlysed cells and large debris. Crude mitochondria were enriched by centrifugation of the supernatant (12,000 g, 15 min, 4°C). The mitochondrial pellet was resuspended in a small amount of cold MIB to a final concentration of 5–10 mg/ml. 200 μg aliquots of mitochondria were flash-frozen in liquid nitrogen after measuring the protein concentration by a Bradford assay (Bio-Rad) and stored at −80°C.

### Alkaline extraction and protease protection analysis

The solubility of Mmc1 was examined by alkaline extraction as described previously^26^, with the following modifications. Equal amounts of crude mitochondria were rinsed with MIB buffer, and the pellets were either immediately resuspended in Laemmli sample buffer or sodium carbonate buffer (0.1 M Na_2_CO_3_, pH 11.0) and incubated for 30 min on ice (mixing every 10 minutes). Mitochondria treated with sodium carbonate buffer were subjected to ultracentrifugation (60 min, 100,000 g, 4°C) and pellets were resuspended in an equivalent volume of Laemmli sample buffer. Sodium carbonate-extracted supernatant proteins were precipitated with TCA (12.5%, 30 min on ice), pelleted by centrifugation (10 min, 16,000 g, 4°C), washed with cold acetone, and resuspended in an equivalent volume of Laemmli sample buffer. Samples were analyzed by SDS-PAGE and Western blotting as described below.

Protease protection analysis of Mmc1 was performed as described previously^26^ with minor modifications. Mitochondria isolated from the strains expressing the indicated chromosomal protein tag were analyzed simultaneously. Crude mitochondria isolated from each strain were thawed, rinsed with MIB, and divided equally into four tubes. After centrifugation, the pellets were resuspended in equivalent volumes of MIB buffer, mitoplast buffer (20 mM HEPES pH 7.4), or solubilization buffer (MIB buffer containing 1% Triton X-100). After 15 minutes of incubation on ice, mitoplast samples were mechanically disrupted by pipetting 15 times. MIB-resuspended mitochondria were mock-treated or proteinase K (100 μg/ml) treated, whereas mitoplast and detergent solubilized samples were treated with proteinase K, and all samples were incubated on ice for an additional 15 min. To inhibit the protease reaction, 2 mM PMSF was added to all samples and incubated for 5 min on ice. Samples were centrifuged (10,400 g, 15min, 4°C) and pellets were resuspended in MIB with 1× protease inhibitor cocktail followed by precipitation with TCA (12.5%, 30 min on ice). Samples were finally resuspended in Laemmli buffer and examined by SDS-PAGE and Western blotting as described below.

### Whole-cell lysate preparation and Western blotting

Whole-cell lysates from *S. pombe* cells were prepared as described^65^ with minor modifications. Briefly, strains were grown exponentially at 30°C in YES media. Cells (5 OD_600_) were pelleted by centrifugation (5000 g for 1 min at room temperature), rinsed with water, and resuspended in 0.3 ml water with a 2× concentration of protease inhibitor cocktail. Cells were diluted 1:2 in a solution of 0.6M NaOH. The samples were incubated for 10 min at room temperature, centrifuged (1000g for 3 min, 4°C), and resuspended in 70 µl of Laemmli buffer prior to Western analysis.

Freshly prepared whole-cell lysates or crude mitochondria extracts were incubated at 95°C for 3 min prior to resolution by SDS-PAGE, transferred to 0.45-μm pore size PVDF membranes, and immunoblotted with the indicated primary antibodies (FLAG (F1804; Sigma-Aldrich), rabbit α-GAPDH (10494-1-AP; Proteintech), mouse α-HA (26183; 1:2000; Thermo Fisher Scientific), rabbit α-GFP (ab290; Abcam). To detect Mic19-3xHA, goat anti-mouse HRP secondary antibody (A4416; Sigma-Aldrich) was used and the signal was observed with SuperSignal West Femto Substrate (Thermo Fisher Scientific). All other proteins were detected with secondary antibodies conjugated to DyLight800 (SA5-35521 and SA5-35571, Thermo Fisher Scientific). A ChemiDoc MP Imaging System (Bio-Rad) was used to capture images and linear adjustments to images were made with Photoshop (Adobe).

### 2D BN-PAGE analysis

2D BN-PAGE analysis was performed as described^27^ with the following modifications. 100 μg of mitochondria isolated from the indicated strains were centrifuged (21,000 × g, 10 min, 4°C), and pellets were resuspended in 20 μl of 1× NativePAGE Sample Buffer (Thermo Fisher Scientific) supplemented with 1× protease inhibitor cocktail and digitonin (SigmaMillipore) to a final detergent:protein ratio of 4 g/g. Samples were solubilized on ice for 15 min followed by centrifugation (21,000×g, 30 min, 4°C). Coomassie Blue G-250 dye was added to the supernatant, which contained solubilized mitochondria, to a final detergent:dye ratio of 16 g/g before running on a 3-12% NativePAGE Mini Protein Gel (Thermo Fisher Scientific) as per manufacturer’s instructions. Protein complex sizes were determined with NativeMark Unstained Protein Standard (ThermoFisher). After resolving the protein complexes under native conditions, the entire lanes were excised and incubated in 10 ml of denaturing buffer (0.12 M Tris-HCl, pH 6.8, 4% SDS, 20% glycerol, and 10% β-mercaptoethanol) for 25 min^66^. Each excised gel slice was microwaved for 10 s halfway through incubation. The gel slices were placed horizontally on a denaturing Tris-Glycine polyacrylamide gel and sealed in position with 0.75% agarose in an SDS-PAGE running buffer, before resolving by SDS-PAGE. Proteins were transferred to 0.45-μm pore size PVDF membranes and analyzed by Western as described above.

### Mammalian cell growth, exogenous Mmc1 expression, and immunofluorescence

U2OS cells (a kind gift of Jodi Nunnari) were grown by culturing in DMEM (Sigma-Aldrich D5796) supplemented with 10% fetal bovine serum (Sigma-Aldrich), 25 mM HEPES, 100 units/ml penicillin, and 100 μg/ml streptomycin.

For expression of Mmc1-GFP and mito-mCherry, U2OS cells were transiently transfected with the plasmids described below using Lipofectamine 3000 (Thermo Fisher Scientific) according to the manufacturer’s instructions. To exogenously express Mmc1, pMmc1-AcGFP-N1 was constructed by PCR amplifying the coding sequence of Mmc1 and cloning into the XhoI/BamHI sites of pAcGFP-N1 by Gibson assembly. mito-mCherry was generated by PCR amplifying the mitochondrial targeting sequence of *N. crassa* Su9 from mito-EGFP^67^ and cloning into the XhoI/BamHI sites of pmCherry-N1 (a kind gift of Gia Voeltz).

For immunofluorescence assays, transfected cells were cultured on glass-bottom cover dishes (CellVis) and were fixed in a 4% paraformaldehyde solution in PBS (15 min, room temperature). Fixed cells were permeabilized (0.1% Triton X-100 in PBS), blocked (10% FBS and 0.1% Triton X-100 in PBS), and then incubated with the indicated primary antibodies (mouse anti- MIC60 [110329; Abcam], rabbit anti-MIC27 [PA5-51427; Thermo Fisher Scientific], mouse anti- TIMM23 [BDB611222; Thermo Fisher Scientific], mouse anti-dsDNA [ab27156; Abcam], rabbit anti-HSP60 [15282-1-AP; Proteintech], or mouse anti-HSP60 [66041-1-Ig; Proteintech]) in blocking solution for 30 min. After incubation, cells were rinsed several times with PBS and incubated with secondary antibodies (donkey anti-rabbit Alexa Fluor 647 Plus [PIA32795; Thermo Fisher Scientific], donkey anti-mouse Alexa Fluor 647 [A-31571; Thermo Fisher Scientific], donkey anti-rabbit Alexa Fluor 555 [A-31572; Thermo Fisher Scientific], and/or donkey anti-mouse Alexa Fluor 555 [A-31570; Thermo Fisher Scientific]) in blocking solution for 30 min. Cells were subsequently rinsed several times with PBS before imaging. Images were acquired on a Nikon Spinning Disk Confocal microscope as described above. Non-transfected control cells were imaged using identical acquisition settings as GFP-transfected cells. ImageJ/Fiji was used to make linear adjustments to images, and linescan analysis was performed as described above.

### Bioinformatic analyses

To search for homologs, the SpMmc1 protein sequence was used as a query in three iterative searches against a database of fungal proteomes from Mycocosm using the jackhammer program of the HMMER v.3.4 suite. The comprehensive phylogenetic tree of Fungi inferred by Li et al. was used as a reference phylogeny^68^. This phylogenetic tree was down-sampled from 1,672 to 477 taxa using Treemmer v.0.3 in such a way that all taxa retained were present in the Mycocosm database and there were no more than three species per genus. Subcellular localization and mitochondrial pre-sequences were predicted with DeepLoc v.2.0. Transmembrane segments were predicted with DeepTMHMM v.1.0.24. Pfam domains were searched against the retrieved sequences using the pHMM’s gathering thresholds with the hmmscan program of the HMMER v.3.4 suite. Proteins were considered Mmc1 candidate homologs if they were predicted to be mitochondrion-localized, had two transmembrane segments towards the C-terminus, and had no other identifiable Pfam domains. Anvi’o v.7 was used to summarize the data associated with the retrieved homologs.

### Statistical analysis

Statistical analysis of fluorescence microscopy was performed as indicated in legends with GraphPad Prism 9.5.1. Comparisons of mitochondrial morphology were performed on the tubular mitochondrial morphology category. Statistical analysis of IP/MS was performed by student’s *t* test using Microsoft Excel.

## Acknowledgments

We thank Madeleine Vaughn for her technical contributions. We thank Mike Henne, Laura Lackner, and Marijn Ford for helpful discussions and Natalie Niemi for the critical review of the manuscript. The UT Southwestern Quantitative Light Microscopy Core Facility, which is supported in part by NIH P30CA142543, provided access to the Nikon spinning disk microscope (purchased with NIH 1S10OD028630-01). We thank Anza Darehshouri and Phoebe Doss from the UT Southwestern Electron Microscopy Core Facility (supported by NIH 1S10OD021685-01A1) for their technical contributions in sample preparation for electron microscopy. We thank the UT Southwestern Proteomics Core Facility for help with mass spectrometry analysis. J.R.F. is supported by a grant from the NIH (R35GM137894) and the UT Southwestern Endowed Scholars Program. M.L.R. is supported by grants from the NIH (R21AI171227 & R01AI150715), Welch foundation (I-2075-20210327), and a Burroughs Wellcome Fund PATH award (1021959).

## References

1. Cogliati, S., Enriquez, J.A., and Scorrano, L. (2016). Mitochondrial Cristae: Where Beauty Meets Functionality. Trends Biochem Sci 41, 261–273. 10.1016/j.tibs.2016.01.001.

2. Klecker, T., and Westermann, B. (2021). Pathways shaping the mitochondrial inner membrane. Open Biol 11, 210238. 10.1098/rsob.210238.

3. Colina-Tenorio, L., Horten, P., Pfanner, N., and Rampelt, H. (2020). Shaping the mitochondrial inner membrane in health and disease. J Intern Med. 10.1111/joim.13031.

4. Mukherjee, I., Ghosh, M., and Meinecke, M. (2021). MICOS and the mitochondrial inner membrane morphology - when things get out of shape. FEBS Lett 595, 1159–1183. 10.1002/1873-3468.14089.

5. Anand, R., Reichert, A.S., and Kondadi, A.K. (2021). Emerging Roles of the MICOS Complex in Cristae Dynamics and Biogenesis. Biology (Basel) 10. 10.3390/biology10070600.

6. Eramo, M.J., Lisnyak, V., Formosa, L.E., and Ryan, M.T. (2020). The ’mitochondrial contact site and cristae organising system’ (MICOS) in health and human disease. J Biochem 167, 243–255. 10.1093/jb/mvz111.

7. Huynen, M.A., Muhlmeister, M., Gotthardt, K., Guerrero-Castillo, S., and Brandt, U. (2016). Evolution and structural organization of the mitochondrial contact site (MICOS) complex and the mitochondrial intermembrane space bridging (MIB) complex. Biochim Biophys Acta 1863, 91–101. 10.1016/j.bbamcr.2015.10.009.

8. Munoz-Gomez, S.A., Slamovits, C.H., Dacks, J.B., Baier, K.A., Spencer, K.D., and Wideman, J.G. (2015). Ancient homology of the mitochondrial contact site and cristae organizing system points to an endosymbiotic origin of mitochondrial cristae. Curr Biol 25, 1489–1495. 10.1016/j.cub.2015.04.006.

9. Rampelt, H., Zerbes, R.M., van der Laan, M., and Pfanner, N. (2017). Role of the mitochondrial contact site and cristae organizing system in membrane architecture and dynamics. Biochim Biophys Acta Mol Cell Res 1864, 737–746. 10.1016/j.bbamcr.2016.05.020.

10. Bohnert, M., Zerbes, R.M., Davies, K.M., Muhleip, A.W., Rampelt, H., Horvath, S.E., Boenke, T., Kram, A., Perschil, I., Veenhuis, M., et al. (2015). Central role of Mic10 in the mitochondrial contact site and cristae organizing system. Cell Metab 21, 747–755. 10.1016/j.cmet.2015.04.007.

11. Barbot, M., Jans, D.C., Schulz, C., Denkert, N., Kroppen, B., Hoppert, M., Jakobs, S., and Meinecke, M. (2015). Mic10 oligomerizes to bend mitochondrial inner membranes at cristae junctions. Cell Metab 21, 756–763. 10.1016/j.cmet.2015.04.006.

12. Bock-Bierbaum, T., Funck, K., Wollweber, F., Lisicki, E., von der Malsburg, K., von der Malsburg, A., Laborenz, J., Noel, J.K., Hessenberger, M., Jungbluth, S., et al. (2022). Structural insights into crista junction formation by the Mic60-Mic19 complex. Sci Adv 8, eabo4946. 10.1126/sciadv.abo4946.

13. Ott, C., Ross, K., Straub, S., Thiede, B., Götz, M., Goosmann, C., Krischke, M., Mueller, M.J., Krohne, G., Rudel, T., and Kozjak-Pavlovic, V. (2012). Sam50 functions in mitochondrial intermembrane space bridging and biogenesis of respiratory complexes. Mol. Cell. Biol. 32, 1173–1188. 10.1128/MCB.06388-11.

14. Xie, J., Marusich, M.F., Souda, P., Whitelegge, J., and Capaldi, R.A. (2007). The mitochondrial inner membrane protein mitofilin exists as a complex with SAM50, metaxins 1 and 2, coiled-coil-helix coiled-coil-helix domain-containing protein 3 and 6 and DnaJC11. FEBS Lett. 581, 3545–3549. 10.1016/j.febslet.2007.06.052.

15. Körner, C., Barrera, M., Dukanovic, J., Eydt, K., Harner, M., Rabl, R., Vogel, F., Rapaport, D., Neupert, W., and Reichert, A.S. (2012). The C-terminal domain of Fcj1 is required for formation of crista junctions and interacts with the TOB/SAM complex in mitochondria. Mol Biol Cell 23, 2143–2155. 10.1091/mbc.E11-10-0831.

16. Zerbes, R.M., Bohnert, M., Stroud, D.A., von der Malsburg, K., Kram, A., Oeljeklaus, S., Warscheid, B., Becker, T., Wiedemann, N., Veenhuis, M., et al. (2012). Role of MINOS in mitochondrial membrane architecture: cristae morphology and outer membrane interactions differentially depend on mitofilin domains. J. Mol. Biol. 422, 183–191. 10.1016/j.jmb.2012.05.004.

17. Friedman, J.R. (2022). Mitochondria from the Outside in: The Relationship Between Inter-Organelle Crosstalk and Mitochondrial Internal Organization. Contact (Thousand Oaks) 5. 10.1177/25152564221133267.

18. Guarani, V., McNeill, E.M., Paulo, J.A., Huttlin, E.L., Frohlich, F., Gygi, S.P., Van Vactor, D., and Harper, J.W. (2015). QIL1 is a novel mitochondrial protein required for MICOS complex stability and cristae morphology. Elife 4. 10.7554/eLife.06265.

19. Rampelt, H., Wollweber, F., Gerke, C., de Boer, R., van der Klei, I.J., Bohnert, M., Pfanner, N., and van der Laan, M. (2018). Assembly of the Mitochondrial Cristae Organizer Mic10 Is Regulated by Mic26-Mic27 Antagonism and Cardiolipin. J Mol Biol 430, 1883–1890. 10.1016/j.jmb.2018.04.037.

20. Anand, R., Kondadi, A.K., Meisterknecht, J., Golombek, M., Nortmann, O., Riedel, J., Peifer-Weiss, L., Brocke-Ahmadinejad, N., Schlutermann, D., Stork, B., et al. (2020). MIC26 and MIC27 cooperate to regulate cardiolipin levels and the landscape of OXPHOS complexes. Life Sci Alliance 3. 10.26508/lsa.202000711.

21. Anand, R., Strecker, V., Urbach, J., Wittig, I., and Reichert, A.S. (2016). Mic13 Is Essential for Formation of Crista Junctions in Mammalian Cells. PLoS One 11, e0160258. 10.1371/journal.pone.0160258.

22. Zerbes, R.M., Hoss, P., Pfanner, N., van der Laan, M., and Bohnert, M. (2016). Distinct Roles of Mic12 and Mic27 in the Mitochondrial Contact Site and Cristae Organizing System. J Mol Biol 428, 1485–1492. 10.1016/j.jmb.2016.02.031.

23. Jimah, J.R., and Hinshaw, J.E. (2019). Structural Insights into the Mechanism of Dynamin Superfamily Proteins. Trends Cell Biol 29, 257–273. 10.1016/j.tcb.2018.11.003.

24. Ford, M.G.J., and Chappie, J.S. (2019). The structural biology of the dynamin-related proteins: New insights into a diverse, multitalented family. Traffic 20, 717–740. 10.1111/tra.12676.

25. Connor, O.M., Matta, S.K., and Friedman, J.R. (2023). Completion of mitochondrial division requires the intermembrane space protein Mdi1/Atg44. J Cell Biol 222. 10.1083/jcb.202303147.

26. Hoppins, S., Collins, S.R., Cassidy-Stone, A., Hummel, E., Devay, R.M., Lackner, L.L., Westermann, B., Schuldiner, M., Weissman, J.S., and Nunnari, J. (2011). A mitochondrial-focused genetic interaction map reveals a scaffold-like complex required for inner membrane organization in mitochondria. The Journal of Cell Biology 195, 323–340. 10.1083/jcb.201107053.

27. Gok, M.O., Connor, O.M., Wang, X., Menezes, C.J., Llamas, C.B., Mishra, P., and Friedman, J.R. (2023). The outer mitochondrial membrane protein TMEM11 demarcates spatially restricted BNIP3/BNIP3L-mediated mitophagy. J Cell Biol 222. 10.1083/jcb.202204021.

28. Stephan, T., Bruser, C., Deckers, M., Steyer, A.M., Balzarotti, F., Barbot, M., Behr, T.S., Heim, G., Hubner, W., Ilgen, P., et al. (2020). MICOS assembly controls mitochondrial inner membrane remodeling and crista junction redistribution to mediate cristae formation. EMBO J 39, e104105. 10.15252/embj.2019104105.

29. Harner, M., Körner, C., Walther, D., Mokranjac, D., Kaesmacher, J., Welsch, U., Griffith, J., Mann, M., Reggiori, F., and Neupert, W. (2011). The mitochondrial contact site complex, a determinant of mitochondrial architecture. EMBO J 30, 4356–4370. 10.1038/emboj.2011.379.

30. von der Malsburg, K., Müller, J.M., Bohnert, M., Oeljeklaus, S., Kwiatkowska, P., Becker, T., Loniewska-Lwowska, A., Wiese, S., Rao, S., Milenkovic, D., et al. (2011). Dual role of mitofilin in mitochondrial membrane organization and protein biogenesis. Dev. Cell 21, 694–707. 10.1016/j.devcel.2011.08.026.

31. Rabl, R., Soubannier, V., Scholz, R., Vogel, F., Mendl, N., Vasiljev-Neumeyer, A., Körner, C., Jagasia, R., Keil, T., Baumeister, W., et al. (2009). Formation of cristae and crista junctions in mitochondria depends on antagonism between Fcj1 and Su e/g. The Journal of Cell Biology 185, 1047–1063. 10.1083/jcb.200811099.

32. John, G.B., Shang, Y., Li, L., Renken, C., Mannella, C.A., Selker, J.M.L., Rangell, L., Bennett, M.J., and Zha, J. (2005). The mitochondrial inner membrane protein mitofilin controls cristae morphology. Mol Biol Cell 16, 1543–1554. 10.1091/mbc.E04-08-0697.

33. Itoh, K., Tamura, Y., Iijima, M., and Sesaki, H. (2013). Effects of Fcj1-Mos1 and mitochondrial division on aggregation of mitochondrial DNA nucleoids and organelle morphology. Mol Biol Cell 24, 1842–1851. 10.1091/mbc.E13-03-0125.

34. Friedman, J.R., Mourier, A., Yamada, J., McCaffery, J.M., and Nunnari, J. (2015). MICOS coordinates with respiratory complexes and lipids to establish mitochondrial inner membrane architecture. Elife 4. 10.7554/eLife.07739.

35. Balsa, E., Soustek, M.S., Thomas, A., Cogliati, S., Garcia-Poyatos, C., Martin-Garcia, E., Jedrychowski, M., Gygi, S.P., Enriquez, J.A., and Puigserver, P. (2019). ER and Nutrient Stress Promote Assembly of Respiratory Chain Supercomplexes through the PERK-eIF2alpha Axis. Mol Cell 74, 877–890 e876. 10.1016/j.molcel.2019.03.031.

36. Ott, C., Dorsch, E., Fraunholz, M., Straub, S., and Kozjak-Pavlovic, V. (2015). Detailed analysis of the human mitochondrial contact site complex indicate a hierarchy of subunits. PLoS One 10, e0120213. 10.1371/journal.pone.0120213.

37. Li, H., Ruan, Y., Zhang, K., Jian, F., Hu, C., Miao, L., Gong, L., Sun, L., Zhang, X., Chen, S., et al. (2016). Mic60/Mitofilin determines MICOS assembly essential for mitochondrial dynamics and mtDNA nucleoid organization. Cell Death Differ 23, 380–392. 10.1038/cdd.2015.102.

38. Jans, D.C., Wurm, C.A., Riedel, D., Wenzel, D., Stagge, F., Deckers, M., Rehling, P., and Jakobs, S. (2013). STED super-resolution microscopy reveals an array of MINOS clusters along human mitochondria. Proc Natl Acad Sci USA 110, 8936–8941. 10.1073/pnas.1301820110.

39. Callegari, S., Muller, T., Schulz, C., Lenz, C., Jans, D.C., Wissel, M., Opazo, F., Rizzoli, S.O., Jakobs, S., Urlaub, H., et al. (2019). A MICOS-TIM22 Association Promotes Carrier Import into Human Mitochondria. J Mol Biol 431, 2835–2851. 10.1016/j.jmb.2019.05.015.

40. Martin-Castellanos, C., Blanco, M., Rozalen, A.E., Perez-Hidalgo, L., Garcia, A.I., Conde, F., Mata, J., Ellermeier, C., Davis, L., San-Segundo, P., et al. (2005). A large-scale screen in S. pombe identifies seven novel genes required for critical meiotic events. Curr Biol 15, 2056–2062. 10.1016/j.cub.2005.10.038.

41. Ikebe, C., Konishi, M., Hirata, D., Matsusaka, T., and Toda, T. (2011). Systematic localization study on novel proteins encoded by meiotically up-regulated ORFs in fission yeast. Biosci Biotechnol Biochem 75, 2364–2370. 10.1271/bbb.110558.

42. Matsuyama, A., Arai, R., Yashiroda, Y., Shirai, A., Kamata, A., Sekido, S., Kobayashi, Y., Hashimoto, A., Hamamoto, M., Hiraoka, Y., et al. (2006). ORFeome cloning and global analysis of protein localization in the fission yeast Schizosaccharomyces pombe. Nat Biotechnol 24, 841–847. 10.1038/nbt1222.

43. Paysan-Lafosse, T., Blum, M., Chuguransky, S., Grego, T., Pinto, B.L., Salazar, G.A., Bileschi, M.L., Bork, P., Bridge, A., Colwell, L., et al. (2023). InterPro in 2022. Nucleic Acids Res 51, D418–D427. 10.1093/nar/gkac993.

44. Tirrell, P.S., Nguyen, K.N., Luby-Phelps, K., and Friedman, J.R. (2020). MICOS subcomplexes assemble independently on the mitochondrial inner membrane in proximity to ER contact sites. J Cell Biol 219. 10.1083/jcb.202003024.

45. Zimmermann, L., Stephens, A., Nam, S.Z., Rau, D., Kubler, J., Lozajic, M., Gabler, F., Soding, J., Lupas, A.N., and Alva, V. (2018). A Completely Reimplemented MPI Bioinformatics Toolkit with a New HHpred Server at its Core. J Mol Biol 430, 2237–2243. 10.1016/j.jmb.2017.12.007.

46. Jumper, J., Evans, R., Pritzel, A., Green, T., Figurnov, M., Ronneberger, O., Tunyasuvunakool, K., Bates, R., Zidek, A., Potapenko, A., et al. (2021). Highly accurate protein structure prediction with AlphaFold. Nature 596, 583–589. 10.1038/s41586-021-03819-2.

47. Varadi, M., Anyango, S., Deshpande, M., Nair, S., Natassia, C., Yordanova, G., Yuan, D., Stroe, O., Wood, G., Laydon, A., et al. (2022). AlphaFold Protein Structure Database: massively expanding the structural coverage of protein-sequence space with high-accuracy models. Nucleic Acids Res 50, D439–D444. 10.1093/nar/gkab1061.

48. Li, Z., Jaroszewski, L., Iyer, M., Sedova, M., and Godzik, A. (2020). FATCAT 2.0: towards a better understanding of the structural diversity of proteins. Nucleic Acids Res 48, W60–W64. 10.1093/nar/gkaa443.

49. Michie, K.A., Boysen, A., Low, H.H., Moller-Jensen, J., and Lowe, J. (2014). LeoA, B and C from enterotoxigenic Escherichia coli (ETEC) are bacterial dynamins. PLoS One 9, e107211. 10.1371/journal.pone.0107211.

50. Wang, M., Guo, X., Yang, X., Zhang, B., Ren, J., Liu, A., Ran, Y., Yan, B., Chen, F., Guddat, L.W., et al. (2019). Mycobacterial dynamin-like protein IniA mediates membrane fission. Nat Commun 10, 3906. 10.1038/s41467-019-11860-z.

51. Quintana-Cabrera, R., and Scorrano, L. (2023). Determinants and outcomes of mitochondrial dynamics. Mol Cell 83, 857–876. 10.1016/j.molcel.2023.02.012.

52. Stiegler, A.L., and Boggon, T.J. (2020). The pseudoGTPase group of pseudoenzymes. FEBS J 287, 4232–4245. 10.1111/febs.15554.

53. Morel, C.A., Asencio, C., Blancard, C., Salin, B., Gontier, E., Duvezin-Caubet, S., Rojo, M., Bringaud, F., and Tetaud, E. (2023). Identification of a novel and ancestral machinery involved in mitochondrial membrane branching in Trypanosoma brucei. bioRxiv. 10.1101/2023.06.28.546890.

54. Vanwalleghem, G., Fontaine, F., Lecordier, L., Tebabi, P., Klewe, K., Nolan, D.P., Yamaryo-Botte, Y., Botte, C., Kremer, A., Burkard, G.S., et al. (2015). Coupling of lysosomal and mitochondrial membrane permeabilization in trypanolysis by APOL1. Nat Commun 6, 8078. 10.1038/ncomms9078.

55. Wang, L., Yan, Z., Vihinen, H., Eriksson, O., Wang, W., Soliymani, R., Lu, Y., Xue, Y., Jokitalo, E., Li, J., and Zhao, H. (2019). FAM92A1 is a BAR domain protein required for mitochondrial ultrastructure and function. J Cell Biol 218, 97–111. 10.1083/jcb.201806191.

56. Bahler, J., Wu, J.Q., Longtine, M.S., Shah, N.G., McKenzie, A., 3rd, Steever, A.B., Wach, A., Philippsen, P., and Pringle, J.R. (1998). Heterologous modules for efficient and versatile PCR-based gene targeting in Schizosaccharomyces pombe. Yeast 14, 943-951. 10.1002/(SICI)1097-0061(199807)14:10<943::AID-YEA292>3.0.CO;2-Y.

57. Longtine, M.S., McKenzie, A., Demarini, D.J., Shah, N.G., Wach, A., Brachat, A., Philippsen, P., and Pringle, J.R. (1998). Additional modules for versatile and economical PCR-based gene deletion and modification in Saccharomyces cerevisiae.

58. Sheff, M.A., and Thorn, K.S. (2004). Optimized cassettes for fluorescent protein tagging in Saccharomyces cerevisiae. Yeast 21, 661–670. 10.1002/yea.1130.

59. Lackner, L.L., Ping, H., Graef, M., Murley, A., and Nunnari, J. (2013). Endoplasmic reticulum-associated mitochondria-cortex tether functions in the distribution and inheritance of mitochondria. Proc Natl Acad Sci USA 110, E458–467. 10.1073/pnas.1215232110.

60. Subramanian, K., Jochem, A., Le Vasseur, M., Lewis, S., Paulson, B.R., Reddy, T.R., Russell, J.D., Coon, J.J., Pagliarini, D.J., and Nunnari, J. (2019). Coenzyme Q biosynthetic proteins assemble in a substrate-dependent manner into domains at ER-mitochondria contacts. J Cell Biol 218, 1353–1369. 10.1083/jcb.201808044.

61. Kraft, L.M., and Lackner, L.L. (2019). A conserved mechanism for mitochondria-dependent dynein anchoring. Mol Biol Cell 30, 691–702. 10.1091/mbc.E18-07-0466.

62. Murray, S. (2008). High pressure freezing and freeze substitution of Schizosaccharomyces pombe and Saccharomyces cerevisiae for TEM. Methods Cell Biol 88, 3–17. 10.1016/S0091-679X(08)00401-9.

63. Gok, M.O., Speer, N.O., Henne, W.M., and Friedman, J.R. (2022). ER-localized phosphatidylethanolamine synthase plays a conserved role in lipid droplet formation. Mol Biol Cell 33, ar11. 10.1091/mbc.E21-11-0558-T.

64. Meeusen, S., McCaffery, J.M., and Nunnari, J. (2004). Mitochondrial fusion intermediates revealed in vitro. Science 305, 1747–1752. 10.1126/science.1100612.

65. Matsuo, Y., Asakawa, K., Toda, T., and Katayama, S. (2006). A rapid method for protein extraction from fission yeast. Biosci Biotechnol Biochem 70, 1992–1994. 10.1271/bbb.60087.

66. Fiala, G.J., Schamel, W.W., and Blumenthal, B. (2011). Blue native polyacrylamide gel electrophoresis (BN-PAGE) for analysis of multiprotein complexes from cellular lysates. J Vis Exp 10.3791/2164.

67. Song, Z., Ghochani, M., McCaffery, J.M., Frey, T.G., and Chan, D.C. (2009). Mitofusins and OPA1 mediate sequential steps in mitochondrial membrane fusion. Mol Biol Cell 20, 3525–3532. 10.1091/mbc.E09-03-0252.

68. Li, Y., Steenwyk, J.L., Chang, Y., Wang, Y., James, T.Y., Stajich, J.E., Spatafora, J.W., Groenewald, M., Dunn, C.W., Hittinger, C.T., et al. (2021). A genome-scale phylogeny of the kingdom Fungi. Curr Biol 31, 1653–1665 e1655. 10.1016/j.cub.2021.01.074.

